# Intragenic methylation repatterning is associated with alternative splicing and unique epigenetic phenotypes

**DOI:** 10.64898/2026.02.18.706621

**Authors:** Alenka Hafner, Hardik Kundariya, Robersy Sanchez, Akshay U Nair, Sally A. Mackenzie

**Affiliations:** Department of Biology, Pennsylvania State University, University Park, Pennsylvania, 16801, United States; Intercollege Graduate Degree Program in Plant Biology, Huck Institutes of the Life Sciences, Pennsylvania State University, University Park, Pennsylvania, 16801, United States; EpiMethyl Analytics, 1005 North Warson Road Ste 401B, St. Louis, MO 63132; Department of Biology, Indian Institute of Science Education and Research (IISER), Tirupati, 517507, Andhra Pradesh, India

## Abstract

The role of intragenic cytosine methylation in shaping phenotypes has been contentious. Recent studies show association between stress and alternative splicing of transcripts, but without functional genome-wide or single-position analysis. We utilized the *msh1* experimental system in Arabidopsis as a model of reproducible epigenetic states with stress-responsive phenotypes, including commitment to heritable memory for at least seven generations. We mapped the methylome to single-cytosine resolution with signal-detection, verified by machine learning. Differentially methylated genes were overlapped with *msh1*-derived transcript isoforms to show that different patterns of exonic methylation led to different levels of isoform expression. Alternatively spliced and differentially methylated genes were enriched in key regulators of growth and development and spliceosome components. Genes targeted for differential methylation also contained a known CTT motif. These results demonstrate a direct relationship in plants between environmentally responsive differential methylation and alternative splicing behavior leading to phenotype changes.

## Introduction

The manner by which environmentally adaptive traits are acquired and inherited remains elusive^1^ despite studies of natural systems demonstrating transgenerational memory^2–8^. DNA methylation of cytosine residues plays a pivotal role in chromatin dynamics, but its influence on phenotypic plasticity remains poorly understood^9,10^. In plants, methylation is found in CG, CHG, and CHH context (where H is a non-guanine nucleotide), and high-density methylation can function in transposable element (TE) silencing and heterochromatin condensation^11^. Low-density, CG methylation generally occurs within gene bodies and is often responsive to environmental change^11^. Amenability of gene body methylation (gbM) to stable inheritance may be important for environmental adaptation^11,12^ but whether it even has a function beyond chromatin stabilization remains controversial^11^.

GbM is often described as simply “bleed through” of aberrant methylation from adjacent promotors and neighboring transposable elements^13–16,11^, but its recent with gene expression variance, independent of TE methylation^12^, goes against that idea. Some evidence seems to show constitutively expressed genes with higher expression levels have more CG methylation but a distinct lack of it at transcription start and termination sites^13,17^. However, this does not preclude gbM from having additional functions inside genes, especially in higher plants with more elaborated machinery able to direct methylation to particular targets^18,19^. Indeed, there is accumulating evidence of selection and gene-regulatory function of intragenic methylation in response to changing environmental conditions and not just in constitutively expressed genes^7,11,12,20–22^.

An emerging hypothesis for how gbM could influence phenotype through gene expression changes, as a requisite for the demonstration of its importance in adaptation, is its impact on alternative splicing. In animal models, small changes in the patterns of gene body methylation have been associated with alternative splicing of transcripts^23–28^. In plants, gbM seems to target genes that correspond to observed phenotypes in environmental response studies^20,29–31^, but overlap between differentially methylated (DMGs) and expressed genes (DEGs) is limited. No analysis platform has offered sufficient resolution in the highly methylated and stochastic plant system to predict methylation effects on phenotype, and association between genic methylation in and alternative splicing has only been described at the genome level or through mutant analysis^11,13,20,32,33^.

The *msh1* system is an experimental model that allows functional study of plant epigenetic responses to environmental stress^21,34–39^ (Fig. 1A). Nuclear gene *MutS Homolog 1* (*MSH1*) encodes a protein targeted to mitochondria and specialized sensory plastids^35,36^, and its downregulation is accompanied by heightened stress responsiveness. The mutants show a range in delayed flowering, dwarfing, and resistance to various environmental stresses^36,40^ (Fig. 1B). Restoration of MSH1 expression by segregating the *MSH1* RNAi transgene triggers persistence of the enhanced stress response for at least seven generations of self-crosses^36,41^. This state of *msh1* ‘memory’ exhibits a uniform, non-genetic phenotype^41^ (Fig. 1B). Use of the *msh1* mutant as rootstock with wild type scion produces graft progeny that exhibit enhanced growth vigor^34^ (Fig. 1B). Deriving these distinct *msh1* states depends on the RNA-directed DNA methylation (RdDM) pathway^34,36,42^ (Fig. 1A).

**Figure 1.**
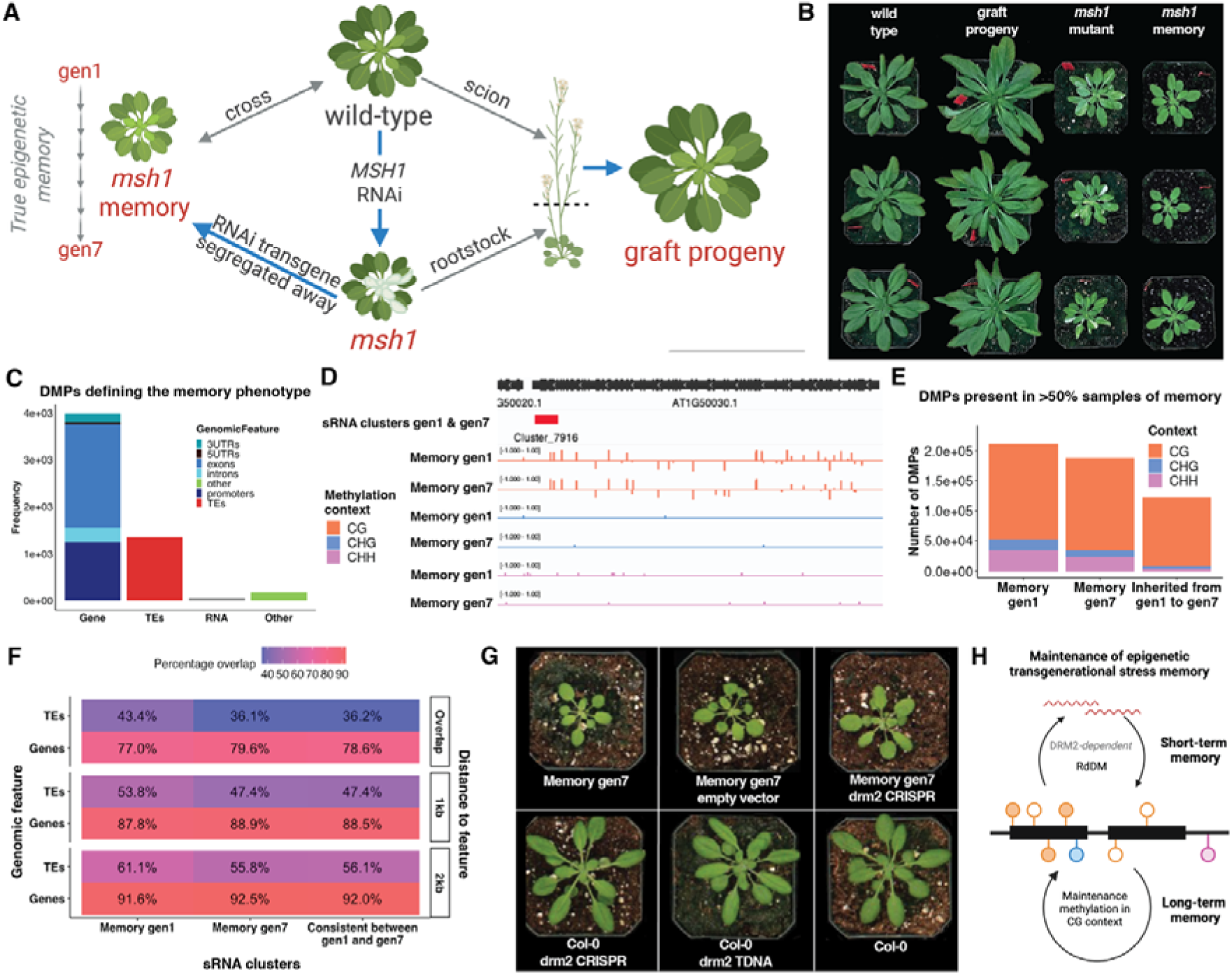
The memory phenotype is inherited for at least seven generations and associated with sRNA and CG methylation. (**A**) The *msh1* experimental model system is composed of distinct epigenetic phenotypes (red).^38^ Generation of *msh1* epigenetic states depends on RdDM machinery (blue arrows) based on mutant analyses^34,41,42^. (**B**) Plant phenotypes (35 days after planting) for wild type (Col-0), progeny of a graft of wild type scion on *msh1* rootstock, *msh1* T-DNA mutants, and *msh1* memory in generation seven. (**C**) Genomic feature distribution of differentially methylated positions (DMPs) that define memory and show inheritance to gen 7. (**D**) A sample genome browser view of memory sRNA clusters (generalized linear regression analysis (GLM), α≤0.05, FDR < 0.05, |log2FC|L≥L0.5) inherited from gen1 to gen7 (red) and CG, CHG, and CHH context DMPs (Hellinger divergence >20% control vs. treatment, GLM, α≤0.05, >95% machine learning classification accuracy) present in gen1 and gen7 at AT1G50030 locus. (**E**) DMPs present in >50% samples, inherited from gen1 and gen7. (**F**) Overlap of sRNA clusters differentially expressed (vs. wild type) in memory gen1, gen7, consistent between gen1 and gen7 with genes or transposable elements (TEs) at distances of ≤2kb, ≤1kb, or ≤0kb (overlap). (**G**) Phenotypes (35 DAP) of *drm2* CRISPR mutants in memory gen7 and wild-type background, with empty vector and T-DNA as controls. (**H**) Model of maintenance epigenetic transgenerational stress memory.

The *msh1* system is a model of natural stress phenotypes with distinct epigenetic states, and is isogenic, unlike the 1001 epigenomes^12^. Hence, it offers a unique opportunity to investigate the relationship between targeted methylation repatterning in gene bodies and alternative spicing, and their potential adjustment of adaptive phenotypes. This system enabled us to study the adaptive adjustment of the whole epigenome, without the limitation of targeted (epi)mutation of one or a small number of genes. Bisulfite sequencing (BS-seq) data were interrogated by a signal detection pipeline that allowed identification of singlet, treatment-associated, differentially methylated positions (DMPs) and their validation through machine learning ^43–45^, allowing higher resolution analysis when compared to standard methylome analysis approaches^43–46^. Features of derived DMGs were then compared to alternatively spliced genes (ASGs) and their differentially expressed isoforms. This approach allowed, for the first time, the identification of sequence and function features of genes targeted for environmentally responsive changes in gene expression via alternative splicing modulated by genic RdDM.

## Results

### Long-term *msh1* stress memory is dependent on CG methylation

In Arabidopsis, introduction of an *MSH1* RNAi construct causes genome-wide changes in cytosine methylation patterns that persist after segregation of the transgene^41^. To investigate what defines the memory phenotype, and whether inter- and trans-generational memory are distinct, deep sequencing (32x) was conducted in generation seven (gen7) of memory and an independent generation one (gen1) memory line, controlling for any stochastic variation between memory lineages. Identification of density-independent, treatment (*msh1*)-associated differential methylation was based on previously described signal detection and machine learning^43–45^ to identify memory-associated changes with highest statistical confidence compared to a previous study^41^ (Supplementary Data 1). Memory in both gen1 and gen7 was characterized by hyper- and hypomethylated DMPs in CG context and hypermethylated DMPs in CHH context, all primarily within TEs and gene bodies (Extended Data Fig. 1). The scale of methylome remodeling, reflected in the total number of DMPs, is consistent with previously published work using signal detection to identify differential methylation^38,39,45,46^ and corresponds to the major phenotype changes between memory and wild type individuals (Fig. 1B) . It was gbM in CG context that showed enrichment, when summed over the total number of cytosines, and stable inheritance for seven generations in two separate memory lineages (Fig. 1C-E).

Initiation of memory is an RdDM-dependent process, and the memory phenotype is not obtained in individuals lacking the methyltransferase *DOMAINS REARRANGED METHYLASE 2* (*DRM2*)^41^. We conducted sRNA sequencing to quantify dependence of memory on RdDM machinery in advanced generations of memory (Supplementary Data 2). Consistent with *DRM2* dependence, gen1 memory contained 10862 unique 21-24bp sRNA clusters when compared to wild type, a scale of change corresponding to the observed scale of methylome and phenotype reprogramming, previously observed inside^42^ and outside of the *msh1* system^19^. Gen7 contained only 2862 unique clusters, of which 2010 sRNA clusters were shared between gen1 and gen7. These clusters predominantly overlapped genes, with 78.6% directly overlapping gene bodies and only 36.2% overlapping TEs (Fig. 1F).

sRNA cluster reduction in gen7 implied that participation of RdDM in retention of memory diminishes over generations. We tested this hypothesis by introducing a *DRM2* CRISPR-Cas9 frameshift mutation at exon 9 in an advanced-generation memory line (Extended Data Fig. 2). We transformed individuals at memory gen4, obtained *drm2* by memory gen7, and compared the phenotype to that of unmodified memory gen7, Col-0 CRISPR *drm2*, Col-0 T-DNA *drm2*, and memory transformed with empty vector (Fig. 1G, Extended Data Fig. 2). Growth rate and juvenility period, both altered features of the *msh1* memory phenotype^41^, were indistinguishable in the memory gen7 *drm2* line from unmodified memory gen7 and wild type (Fig. 1G, Extended Data Fig. 2). Flowering time in memory *drm2* lines shifted from late, as in memory, to early, as in Col-0 *drm2* mutants, with wild type intermediate in timing (Extended Data Fig. 2). These observations indicate that while initiation of memory required RdDM, this dependence diminished by gen7, evidenced by sharply reduced gen7 sRNA cluster numbers and persistence of memory phenotype following *drm2* mutation (Fig. 1H).

### Most differentially methylated genes in memory plants are alternatively spliced

Successful association of heritable phenotype with CG methylation for at least seven generations allowed us to investigate the connection between differential methylation and gene expression. We used long-read RNAseq (Iso-seq) to identify alternatively spliced transcripts in leaf and flower tissues of *msh1* mutant, memory, and graft progeny plants when compared to wild type with EdgeR and FLAIR pipelines (Supplementary Data 3), and we analyzed the overlap of gene expression features with methylome repatterning. Looking at genes with 7-generation memory DMPs in gene bodies, promoters, or within 2kb of a containing TE, we found that integration of differential gene expression^41^, tissue-specific translatome^21^, and alternative splicing datasets accounted for 71.0% (Fig. 2A). In all cases, alternative splicing accounted for the largest share of differential expression, particularly for genic methylation changes (Extended Data Fig. 3). DMPs inherited to gen7 and present in alternatively spliced genes were primarily within exons (Fig. 2B).

**Figure 2.**
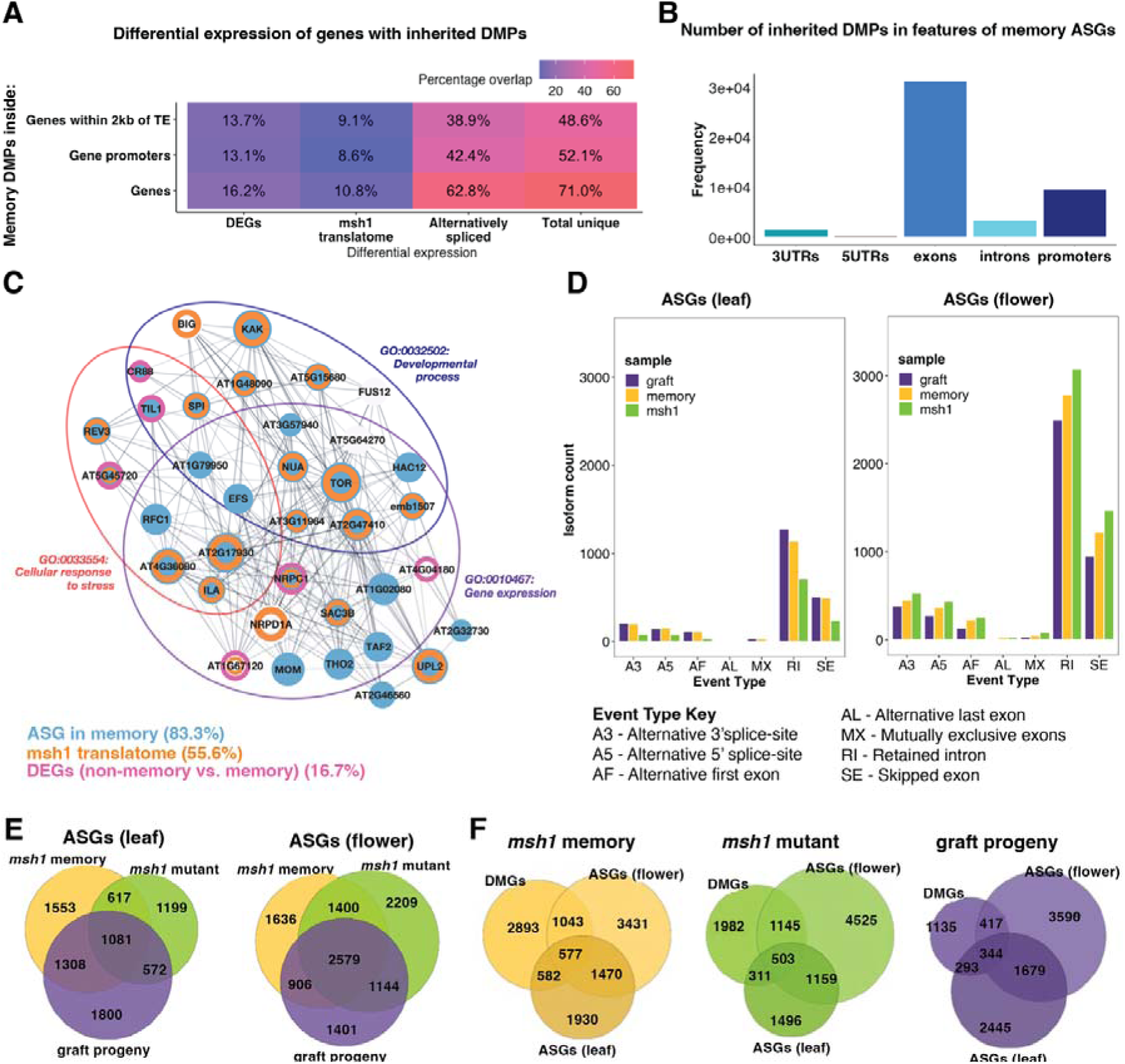
Differentially methylated genes in *msh1* epigenetic phenotypes are alternatively spliced. (**A**) Overlap of differentially expressed genes, as identified by RNA-seq (DEGs), tissue-specific translation in *msh1* (translatome obtained by TRAP-seq), and alternatively spliced genes with DMPs defining memory and inherited to gen7. (**B**) Distribution of DMPs inherited from gen1 to gen7 within alternatively spliced genes in gene body features. (**C**) Core PPI network of 36 hub DMGs defining the memory phenotype. The hub genes belonging to enriched GO terms are circled (FDR < 0.05) and colored according to their differential expression. (**D**) Frequency and type of alternative splicing events in leaf and flower alternatively spliced genes (ASGs) (|log2FC|L≥L0.5 and p-value < 0.05) present in *msh1* mutant, memory and graft progeny states as identified by Iso-seq. (**E**) Venn diagrams showing the overlap of the three *msh1* epigenetic states based on ASGs in leaf and flower tissues. (**F**) Venn diagrams showing the overlap of leaf and flower ASGs with DMGs in the three *msh1* epigenetic states.

The DMGs that defined memory, when compared to their full siblings with a wild type phenotype, formed a network hub of 36 genes through PPI gene network analysis with k-means clustering^39,42,45^ in Cytoscape (Fig. 2C). The derived core hub network contained genes involved in stress response, gene expression, and development, consistent with the observed memory phenotype (Fig. 2C). Alternative splicing, differential gene expression, and *msh1* translatome accounted for altered expression of 34/36 core hub DMGs, with 30/36 alternatively spliced (Fig. 2C). All 36 of the genes overlapped with PolIV-dependent sRNA clusters, as defined by Zhou et al.^19^, and were confirmed *dcl2/3/4* targets according to Kundariya et al.^34^. Only 13/36 were located within 2kb of a TE (Extended Data Fig. 4). Several of the hub genes function in RdDM, including the large subunit of RNA polymerase IV, *NRPD1A*^47^. 30/36 network hub genes directly overlapped with gen1 memory sRNA clusters, whereas 13/36 overlapped with gen7 clusters, and 9/36 overlapped an inherited sRNA cluster (Extended Data Fig. 4), consistent with the diminishment of RdDM effect over generational time in memory. Our ability to detect DMGs that define memory phenotype and significantly overlap changes in gene expression, predominantly by alternative splicing, supports a relationship between these processes.

### Differentially methylated genes are associated with distinct alternative splicing behavior in *msh1* states

We demonstrated in previous studies that different *msh1* states have distinct, reproducible methylome signatures in genes associated with their observed phenotype^34,38,41^. However, the overlap between DMGs and DEGs (identified by RNA-seq) was poor, similar to studies by others, leaving open the question of whether genic methylation repatterning impacts phenotype. To test the relationship of methylome repatterning with tissue-specific expression of alternative isoforms, we analyzed ASGs and DMGs in all three *msh1* states. We quantified splicing events leading to these isoforms with the SUPPA package (Fig. 2D) and identified DMGs present in the three states with an advanced version of the MethylIT signal-detection pipeline, providing higher resolution compared to Kundariya *et al.*^38^ (Supplementary Data 1).

There was significant overlap in ASGs, both flower and leaf, with DMGs shared among all three *msh1* states (Fig. 2E, F). The same was true for DMGs, which were also alternatively spliced (AS-DMGs) (Extended Data Fig. 5). While the distinct *msh1* states shared significant numbers of ASGs, DMGs, and AS-DMGs, they also showed unique sets of genes with differential methylation and alternative splicing that corresponded to their divergent phenotypes (Fig. 2F, Extended Data Fig. 5). Most alternative splicing events were associated with exon skipping and intron retention (Fig. 2D). Unique isoforms were shared amongst at least two of the three *msh1* states, but with different expression levels (Supplementary Data 3). From these comparisons we concluded that overlapping groups of genes served as targets of differential methylation in all three states, a significant portion alternatively spliced. The distinct phenotypes are associated with differentially expressed isoforms, suggesting a potential link between isoform changes and phenotype differences.

### Alternatively spliced, differentially methylated genes appear to be core regulators of *msh1*-derived phenotypes

Because memory DMGs were uniquely enriched in genes for environmental signaling, chromatin remodeling, and gene expression, and AS-DMGs were enriched in regulators of growth and development (Extended Data Fig. 6), we investigated this pattern for plausible directionality. PPI gene network analysis in Cytoscape, based on common DMGs between the three *msh1* states, identified a core network of 126 genes by k-means clustering (Fig. 3A, Supplementary Results, Extended Data Fig. 7). Of these, 21% (26) were alternatively spliced in all three *msh1* states in at least one tissue, mostly flowers (Fig. 3A). 86% (108) of the DMGs were alternatively spliced in at least one state, with different AS-DMG sets in each state.

**Figure 3.**
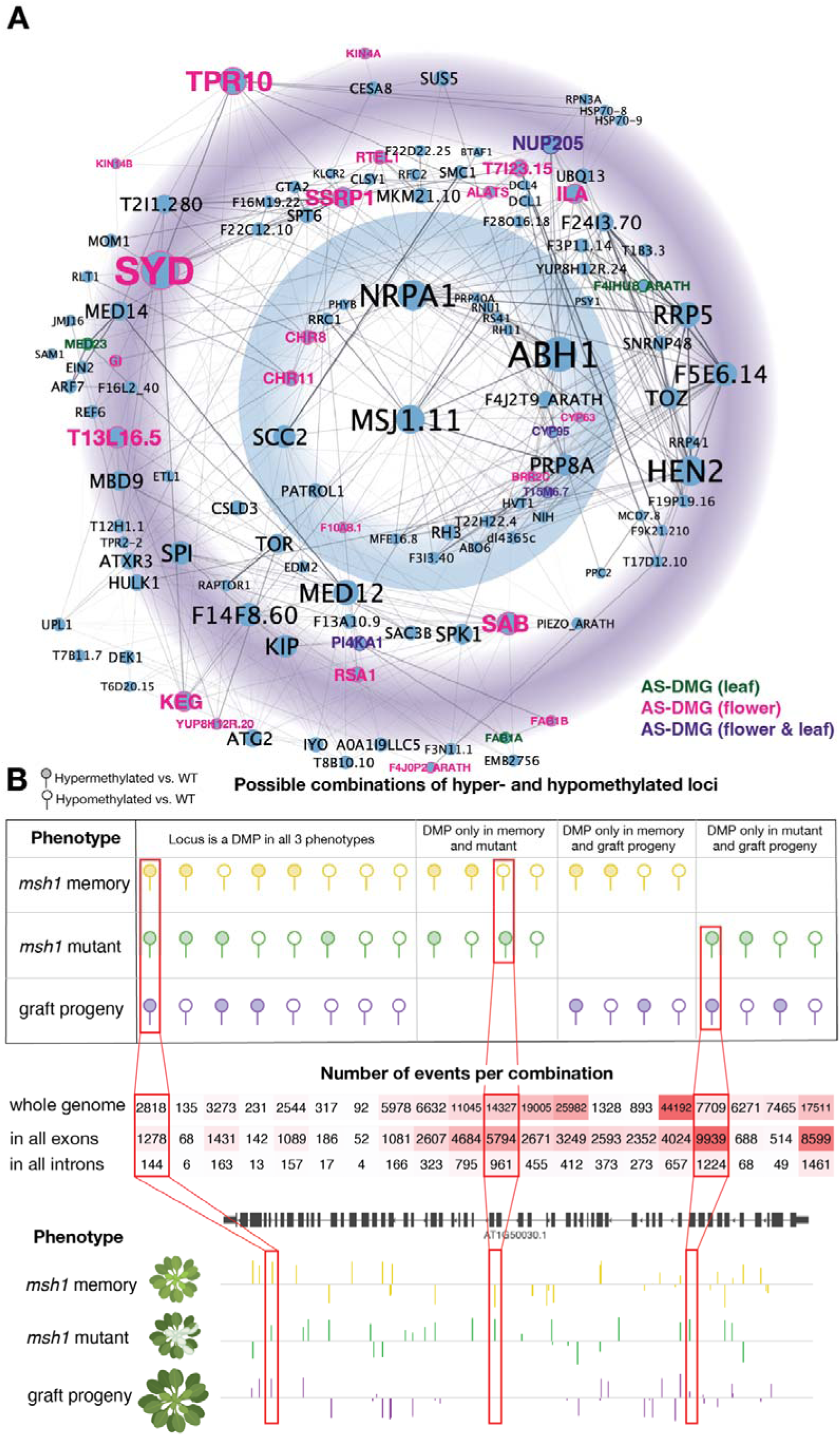
The *msh1* epigenetic phenotypes are defined by a network of alternatively spliced genes with distinct intragenic methylation patterns. (**A**) Core PPI network of 126 DMGs obtained using k-means clustering in Cytoscape from DMGs common between all three *msh1* states. Using the yFiles radial layout algorithm, the DMG nodes were ordered in concentric circles based on their centrality. All genes are DMGs (blue-filled nodes). The blue concentric circle denotes central regulators of the network with the highest level of centrality (network radius <2) and the purple circle denotes proteins with lower centrality (network radius >2). Gene name colors denote DMGs that are also ASGs (pink for flower tissue, green for leaf tissue and purple for both tissues). Most AS-DMGs reside in the outer circle of centrality, indicating that DMGs lie upstream of ASGs. (**B**) *msh1* states are defined by unique combinations of hyper- and hypomethylated loci. The upper panel shows a lollipop diagram of all possible combinations of hyper- and hypomethylation for each cytosine in the genome in the three *msh1* states, excluding loci with a differentially methylated position in only one state. The table contains the counts of all instances of each combination identified as a DMP (total variation>20% WT vs. *msh1* state, GLM, α≤0.05, >95% machine learning classification accuracy). The genome browser view is of all DMPs present in each of the *msh1* states at the AT1G50030 locus with example DMP combinations boxed in red and related to corresponding sections in the lollipop diagram and table.

The genes were arranged radially based on their centrality parameters as measures of their importance in the network beyond their degree of connectivity^48^. Genes with highest centrality (Fig. 3A, inner blue circle) and degree (Fig. 3A, size of node is proportional to degree) were DMGs, but not alternatively spliced in all states (e.g. *NRPA1*, *ABH1*). Of these central genes, 23/28 were alternatively spliced in at least one *msh1* state. On the other hand, genes with lower centrality but high degree were AS-DMGs in all three states, mostly in flower tissue (e.g. *SYD, TPR10*). The group of 126 genes were identified as candidate regulators of the transition between the three *msh1* phenotypes, so we sought to further annotate their functions (Supplementary Data 4, Table 1). ASGs and DMGs were also enriched in regulation by phosphorylation (Supplemental Results, Supplementary Data 5). Together, the core hub network of DMGs in all three *msh1* states supported a hierarchical model in the relationship between gene methylation and subsequent changes in gene expression, with unique sets of DMGs alternatively spliced in different *msh1*-derived phenotypes. We, therefore, postulate that the 126 core hub network genes represent regulators of the epigenetic phenotypes that differentiate the *msh1* states.

**Table 1.**
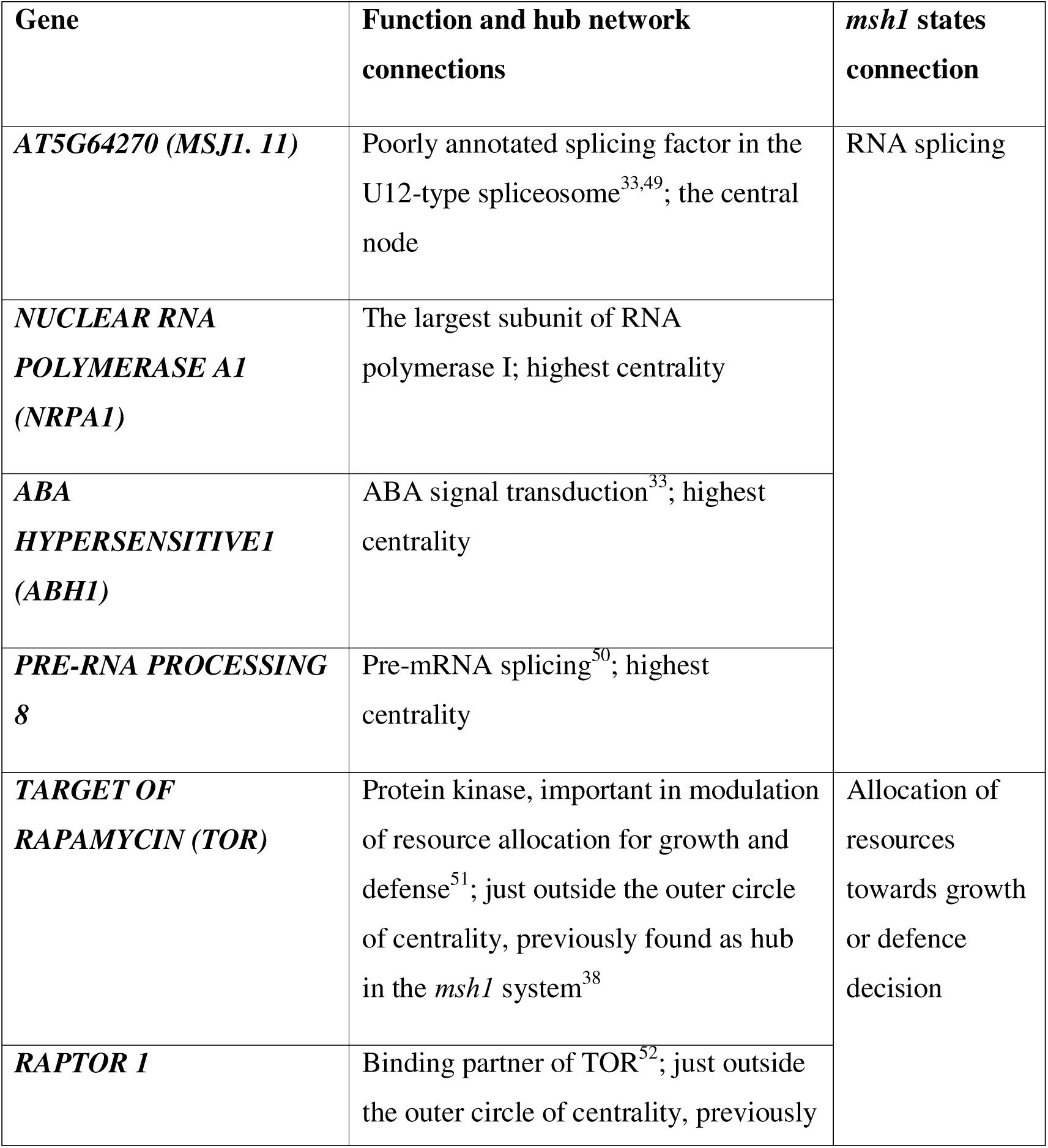
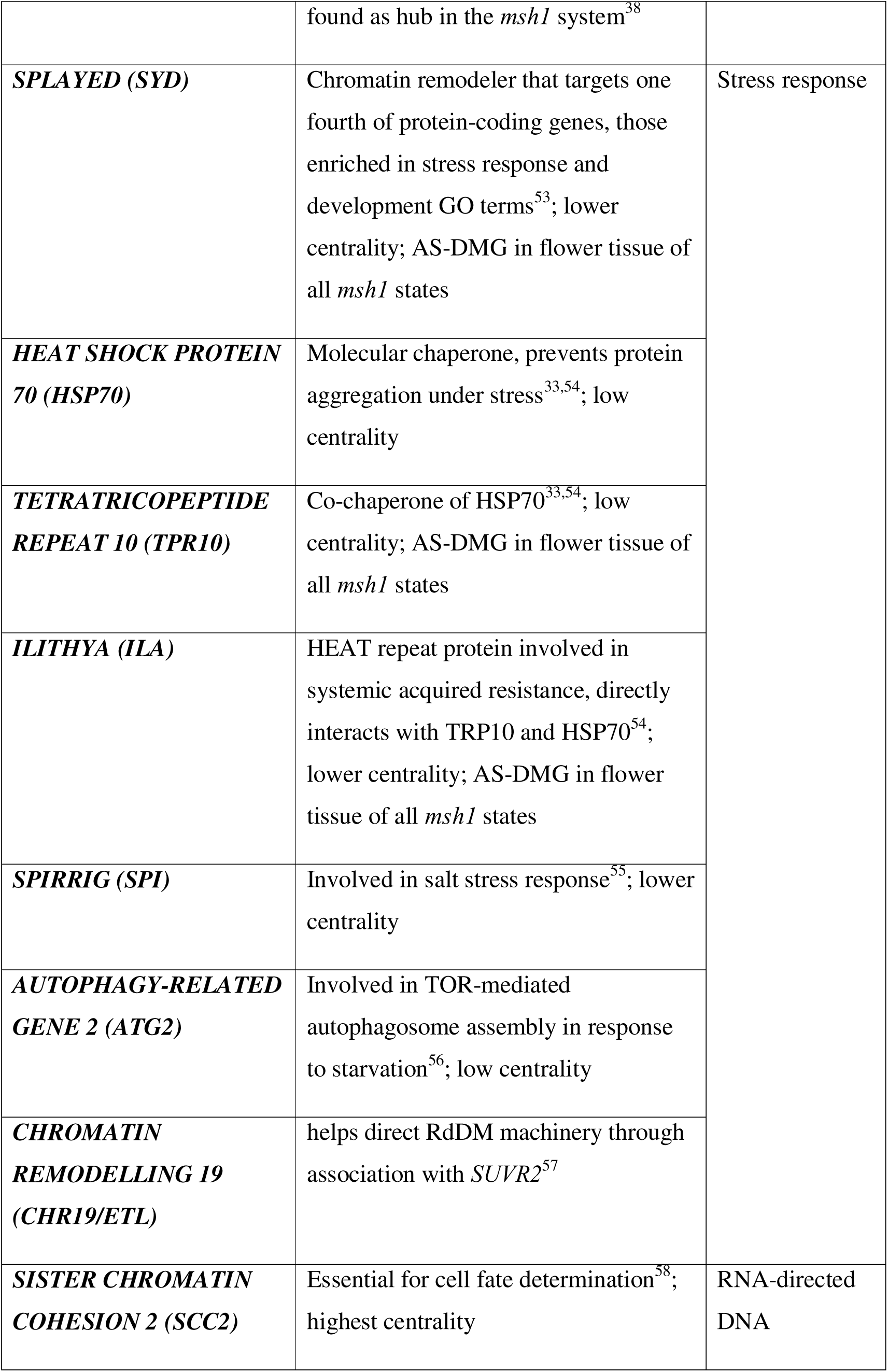
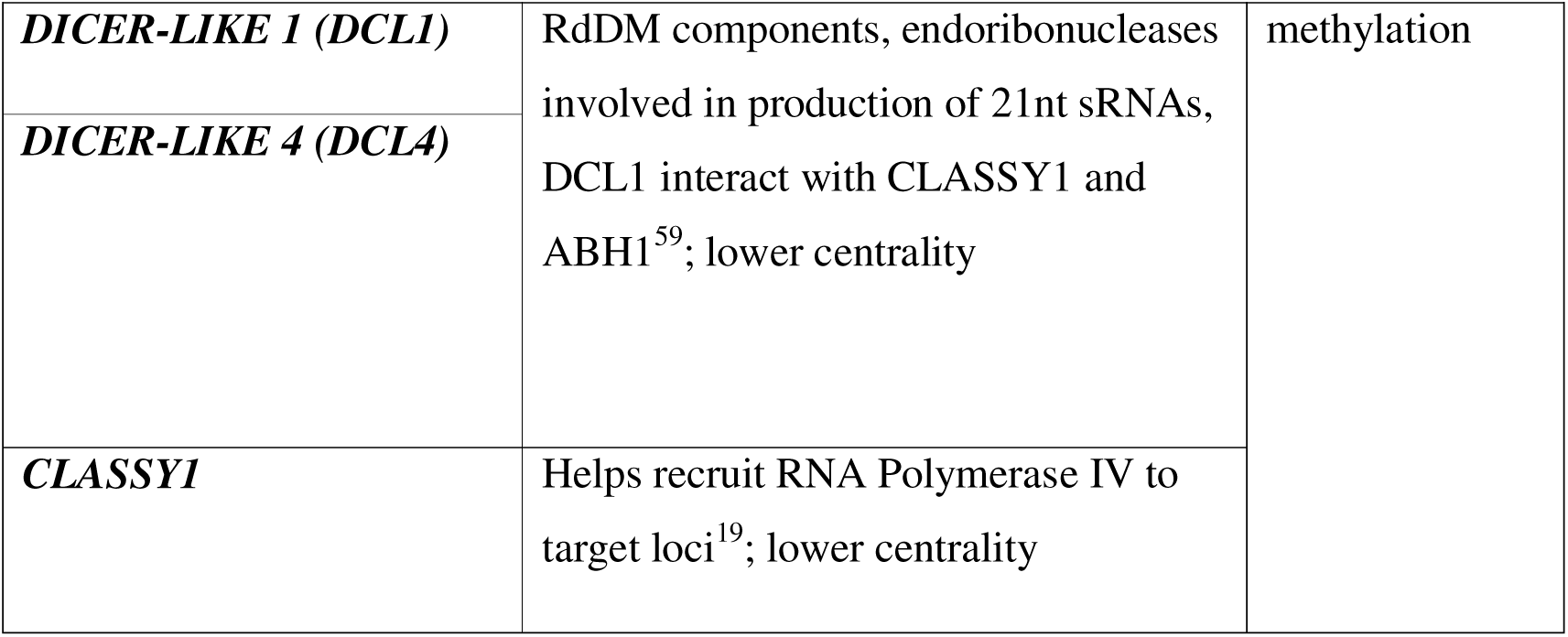
Sample DMGs from the *msh1* states core PPI 126 gene hub network (Fig. 3A) and their connection to *msh1* phenotypes.

### The *msh1* states are distinguished by direction and position of differential methylation

The three *msh1* states shared 1077 DMGs, with another 2045 shared by at least two states (Extended Data Fig. 5). To investigate the relationship of these gene sets with distinct phenotypes, we quantified the instance of all possible combinations of hyper- and hypomethylated DMPs, relative to wild-type, among *msh1* states (Fig. 3B). By comparing frequencies of each DMP combination at whole genome and intragenic levels, variation within exons could account for most state-specific DMP patterns (Fig. 3B), as was the case for detailed analysis of memory (Fig. 1C). These observations support our hypothesis that patterns of individual DMPs and their methylation status within exons provide sufficient information to distinguish *msh1* states and their formation (a direct vs. transgenerational link to *MSH1* suppression). These data serve to increase resolution of dynamic methylome features, which previously pointed generally to genic regions^38^.

### An exonic DNA motif is correlated with RdDM-dependent alternatively spliced isoforms in *msh1* phenotypes

Small RNA clusters can be targeted to particular DNA motifs^19^, and the memory sRNA clusters in our study were targeted to at least 2010 loci consistently for seven generations. We, therefore, investigated sequence similarity among the 36 hub DMGs that define memory. Sequence clustering analysis with Multiple Em for Motif Elicitation (MEME) revealed an enriched 21-bp CTT repeat motif present 180 times in the 36 hub genes (Fig. 4A). A similar, repeating CTT motif could be identified in the *msh1*-associated 67 core hub genes reported previously^38^ (Extended Data Fig. 8A). This 21-bp CTT repeat motif occurred at 98,043 sites in the Arabidopsis genome, most often in exons (Fig. 4B).

**Fig. 4.**
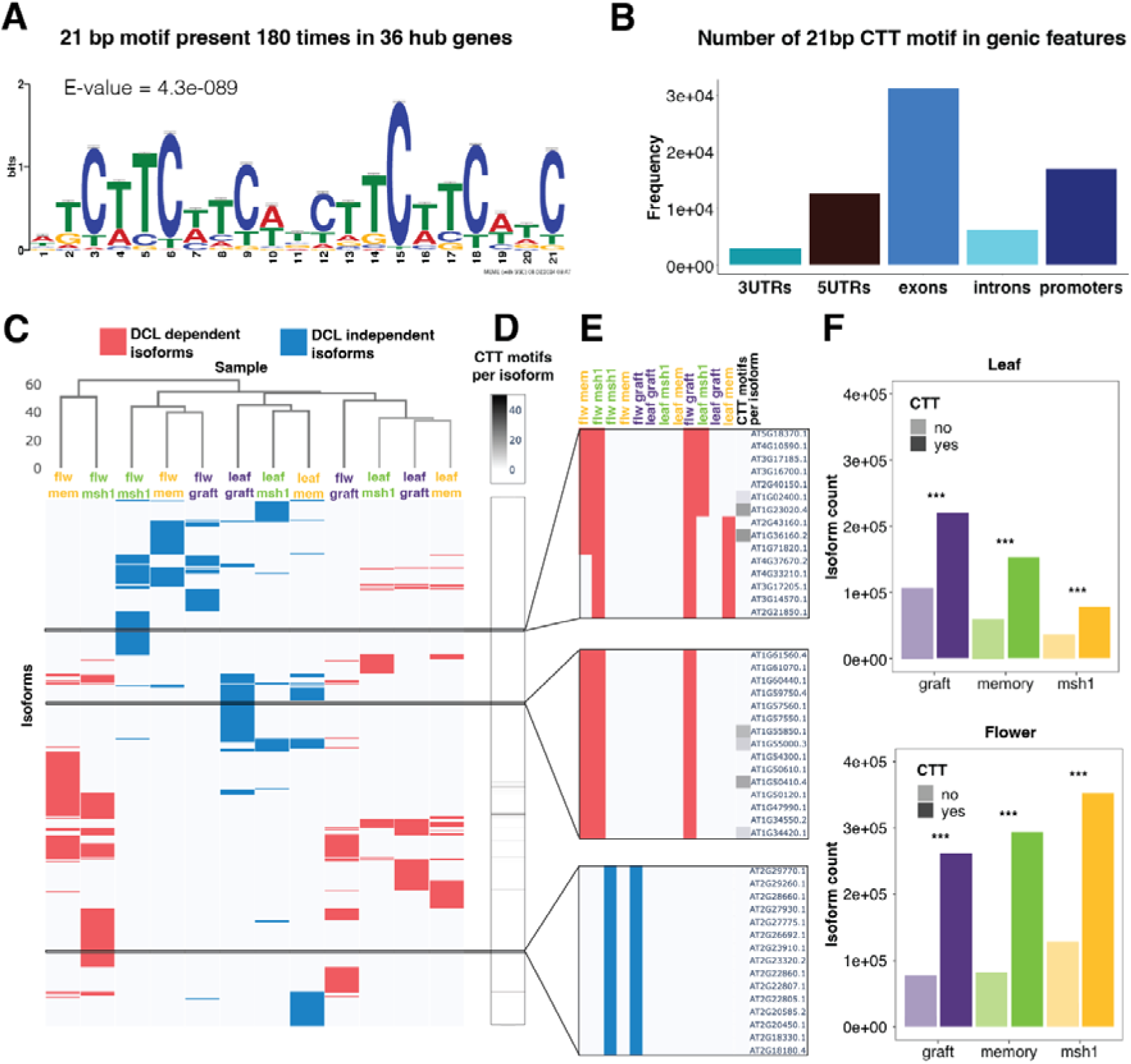
Differentially expressed isoforms that are DCL-dependent can define *msh1* states and coincide with a DNA motif. (**A**) Logo plot of 21-bp motif identified within sequences of 36 core hub DMGs using MEME (ungapped, 6-50 bp, any number of repetitions) (**B**) Distribution of 21-bp CTT motif within genes. (**C**) Hierarchical clustering of differentially expressed isoforms (|log2FC|L≥L0.5 and p-value < 0.05) in *msh1* states for leaf and flower tissue using Ward linkage. *DCL2/3/4*-dependent (red) and independent (blue) isoforms were identified as differentially expressed in *msh1* vs. *msh1dcl2/3/4* and clustered separately. (**D**) The presence and count of the 21bp CTT motif in isoforms clustered in (B). (**E**) Three zoom-in windows of 15 isoforms, their DCL-dependence and presence of CTT motif. (**F**) The absolute number of expressed isoforms in each *msh1* state in leaf and flower tissue and whether each overlaps the 21bp CTT motif (Pearson’s chi-squared, p-values < 0.001).

Because *msh1* state-specific methylome reprogramming relies on RdDM components, we can assume the same of *msh1*-associated alternative splicing. Therefore, we tested for a causal relationship between methylome reprogramming and alternative splicing in *msh1* states as a distinct set of RdDM-dependent isoforms. We used the triple mutant *DICER-LIKE 2, 3,* and *4* (*dcl2/3/4*) in comparisons of *msh1* with *msh1dcl2,3,4* to discriminate DCL*-*dependent isoforms in relation to methylome repatterning (Extended Data Fig. 9). With these data, we constructed a Boolean matrix of all alternatively spliced isoforms, their presence in each *msh1* state for each sampled tissue, and their DCL-dependence. Using hierarchical clustering with Ward linkage, we found that grouping by DCL-dependence superseded grouping by tissue or state (Fig. 4C). Excluding DCL-dependent isoforms found in graft progeny, the clustering of isoforms also produced grouping by tissue.

From derived data, we speculated that the 21bp CTT motif could serve as a putative RdDM target in the *msh1* system, given its prevalence in memory hub genes. However, the CTT codon encodes serine, and serine-rich repeats are key features of the SR family of proteins regulating RNA splicing homeostasis^60^. For example, 33 (26%) of the 126 network genes are direct targets of the well-studied SR45 splicing factor that targets an RNA repeat motif rich in arginine-serine repeats^61^. Mapping all 21bp CTT motifs onto the isoform heatmap (Fig. 4C; Extended Data Fig. 9) showed that 45% (4685) of the 10312 identified isoforms contained one CTT motif (Fig. 4D), and almost 1% (94) contained more than one 21bp CTT motif (Fig. S3B). Of the 4685 CTT-containing isoforms, 22% (1045) were in targets of the SR45 splicing factor, and of the 94 genes with more than one 21bp motif, 26% (24) were in targets of SR45^61^. Browsing 15 sample isoforms suggested a connection between DCL-dependence and the CTT motif (Fig. 4E).

Approximately a quarter of all isoforms appeared DCL-dependent in all states (Supplementary Data 3). Absolute count of expressed transcript isoforms revealed that most isoforms contained a CTT motif (Fig. 4F), with the vast majority DCL-independent (Extended Data Fig. 9B). However, when normalized by gene length to counteract the correlation between gene length and transcript number^62^, CTT motif number per bp depended on DCL presence and not on the number of transcripts produced from the gene (Extended Data Fig. 9C). Furthermore, the relative ratio of isoforms overlapping a CTT motif versus isoforms without was more than 3x higher for DCL-dependent ASGs in all states (Extended Data Fig. 9C). These data support association of the CTT motif with RdDM targets that are alternatively spliced.

### *msh1* target genes are defined by methylation repatterning, CTT motifs and alternative splicing behavior

Isoforms of the putative regulators of *msh1* phenotypes (Fig. 3A), differentially expressed in each state, corresponded to observed phenotypes (Table 2). For example, *SYD* is an AS-DMG in flower tissue of all three states. The gene contained several CTT motif clusters and a PolIV-dependent sRNA cluster present in memory gen1 (Fig. 5A). Full-length and a truncated form (*SYD*Δ*C*) of the gene appear in the nucleus, with the N-terminal domain sufficient for its biological activity^63^. A previous study unable to detect *SYD*Δ*C* mRNA concluded that it was likely a result of post-transcriptional modification^63^. However, long-read sequencing showed *SYD*Δ*C* in flower tissues of all three *msh1* states when compared to wild type, with the full-length form downregulated and *SYD*Δ*C* upregulated in graft progeny (Fig. 5B, C). The upregulation of *SYD*Δ*C* relative to full-length isoform is associated with increased growth vigor^63^, which corresponds to the graft progeny phenotype.

**Figure 5.**
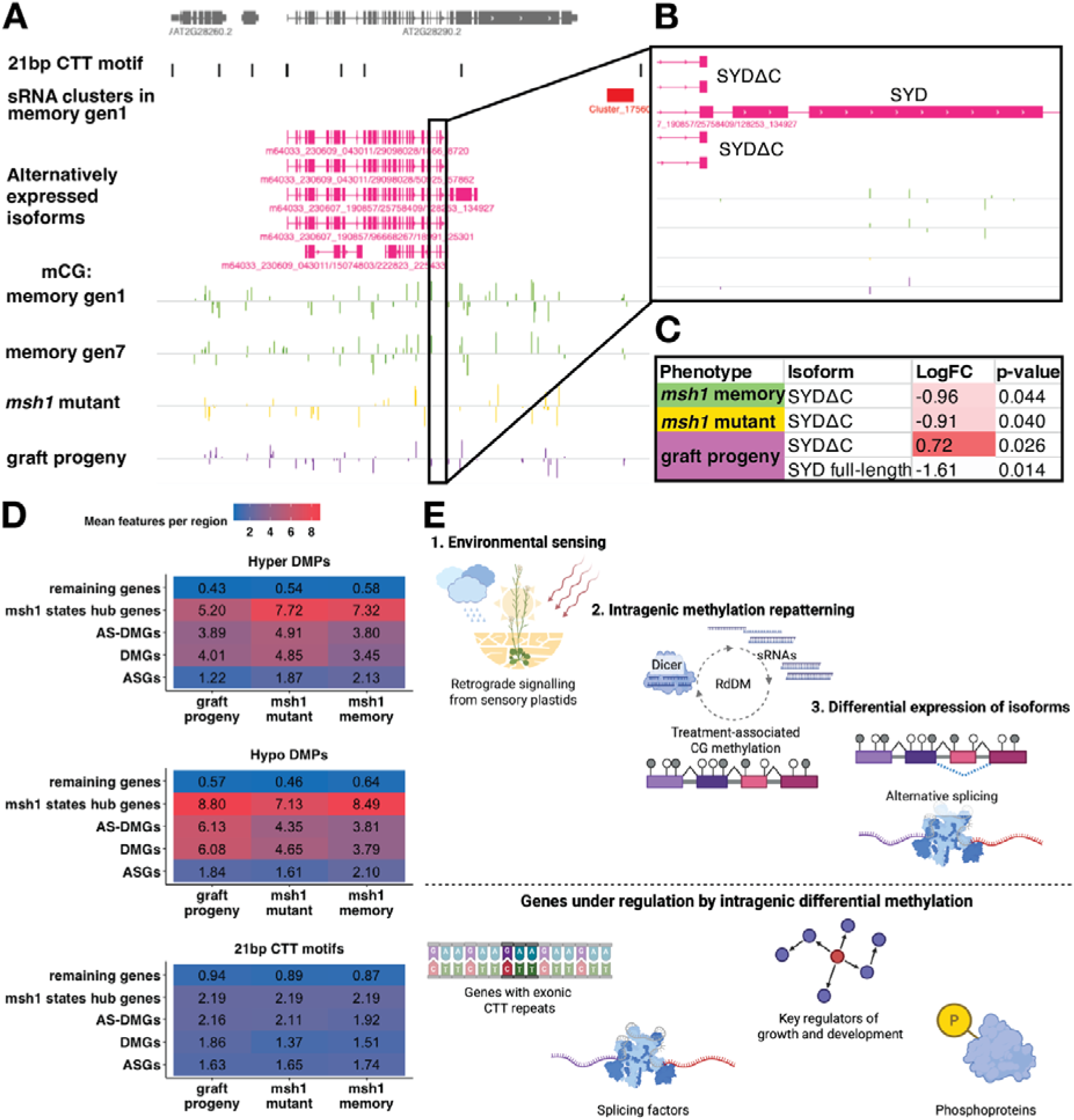
Differentially expressed isoforms define *msh1* states and coincide with a DNA motif when DICER-LIKE (DCL)-dependent. (**A**) A genome browser view of the AT2G28290 (*SPLAYED*) locus with location of 21bp CTT motifs (black), memory gen 1 sRNA clusters (generalized linear regression analysis (GLM), α≤0.05, FDR < 0.05, |log2FC|L≥L0.5) (red), alternatively spliced isoforms present in *msh1* states compared to wild-type (pink), CG context differentially methylated positions (DMPs) (Hellinger divergence >20% control vs. treatment, GLM, α≤0.05, >95% machine learning classification accuracy) present in memory gen1 and gen7 (green), *msh1* mutant (yellow), and graft progeny (purple). (**B**) Zoomed-in view of differentially expressed isoforms of full-length SYD and truncated SYDΔC (**C**) and its expression level in *msh1* states as identified via Iso-seq using EdgeR compared to wild-type tissue (|log2FC|L≥L0.5, p-value < 0.05). (**D**) Mean number of epigenetic features per region of interest in each *msh1* state. The number of hyper-, hypomethylated differentially methylated positions (DMPs), and CTT motifs in each region of interest were divided by the total count of each region of interest. The regions of interest are alternatively spliced genes (ASGs), differentially methylated genes (DMGs), alternatively spliced differentially methylated genes (AS-DMGs), the core hub 126 genes of the *msh1* states derived using protein-protein interaction network analysis and k-means clustering, and all remaining genes in the genome (not ASGs, DMGs, or AS-DMGs). (**E**) Graphical model of gene expression changes modulated by intragenic methylation changes in response to environmental conditions. Following environmental sensing by sensory plastids, a subset of genes is targeted for RNA-directed DNA methylation (RdDM), leading to differential expression of isoforms (upper panel). These environmentally responsive genes represent key regulators of growth and development, are involved in alternative splicing, and are more likely to be phosphoproteins (lower panel).

**Table 2.**
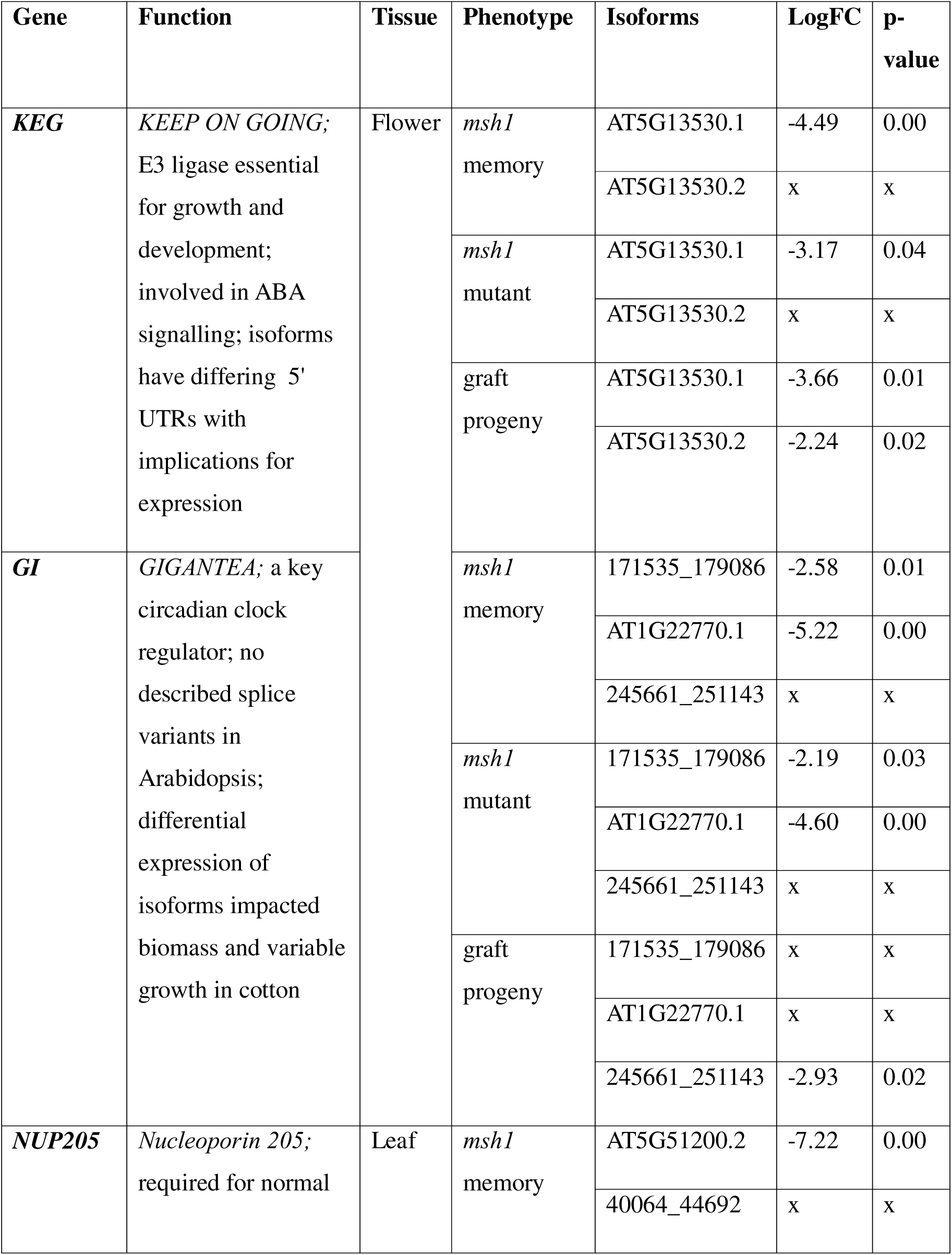
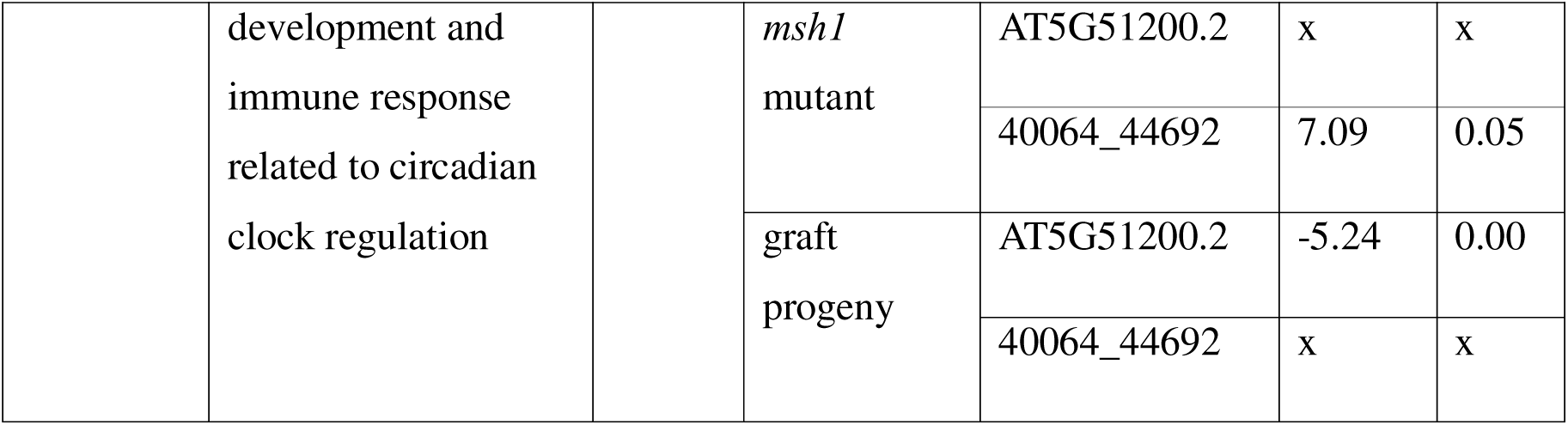
Expression of sample gene isoforms from Fig. 3A, identified via Iso-seq using EdgeR compared to wild type tissue (|log2FC| ≥ 0.5, p-value < 0.05), and their functions. Gene isoforms present in at least one form are listed with their LogFC and an ‘x’ when the isoform is not differentially expressed. ^64–69^

To more fully assess epigenetic and sequence features of DMGs, ASGs, and AS-DMGs identified in this study, we calculated the mean number of hypo- and hypermethylated DMPs and CTT motifs contained in each of the three *msh1* states (Fig. 5D). Background levels of hypo- and hyper-DMPs and CTT motifs in the genome fell below 1.0 across genes not differentially methylated or alternatively spliced. ASGs contained, on average, 1.7 CTT motifs and at least one DMP of each methylation direction (Fig. 5D). DMGs, as defined in our study, contain more than three DMPs of each direction and a similar number of CTT motifs as ASGs (Fig. 5D). AS-DMGs contained the same average number of DMPs but contained, on average, two CTT motifs (Fig. 5D). Two CTT motifs were also found in the 126-hub network of *msh1* states, which contained almost twice as many DMPs in either direction (Fig. 5D). Hence, enrichment of the CTT motif in AS-DMGs further supported its role in their establishment, while DMP density further established the 126-hub genes as targets of *msh1* epigenetic remodeling.

## Discussion

Classically defined gene-body methylation is paradoxical; it comprises exonic DNA methylation in CG context that can be stably transmitted across generations, but has yet to show a role in shaping environmentally-responsive phenotype^11,70^. Alternative splicing regulation has been proposed as a function of gene body methylation in plants, but has lacked the substantial supporting evidence of mammalian systems^11,13,26,70^. With *msh1* memory state as model, we have uncovered a connection between differential methylation and alternative splicing, extending our analysis to three distinct epigenetic phenotypes (states).

Overlap between DMGs and ASGs was significant in all three *msh1* states (Fig. 2). These results match findings from plants under different stress conditions^20,32,33,71^, supporting the assertion that methylation repatterning followed by differential isoform expression is part of plant stress response (Fig. 5E). Approximately a third of Arabidopsis genes are affected in at least one *msh1* state, but the majority of these changes involve stoichiometric adjustment of alternative isoforms, with total transcript count per gene remaining almost unchanged. The purely epigenetic shifts between *msh1* phenotypes suggest that this effect is sufficient to adjust plant form and development. Our observations also support a role of gbM in the adaptive regulation of alternative splicing, not just passive prevention of intron retention^11^. This alternative model would explain contrary evidence from the study of *met1* epiRILs^13^, which were not exposed to changes in environment.

A CTT motif, which associates with intergenic recombination ^72–76^, is also a known target of the SR45 splicing factor, at RNA level, and correlates with targeting of memory sRNAs in our study (Fig. 4). CTT motif number was unrelated to the likelihood of a gene being alternatively spliced, and it is possible that longer CTT repeats are responsible for recruiting RdDM machinery while shorter CTT motifs are sufficient to promote alternative splicing at these sites. While the precise mechanism underlying this association remains unclear, our observations suggest several plausible routes of interaction, which warrant further investigation in future studies. RdDM tissue specificity is partially directed by a distinct DNA motif identified in a CLASSY3-dependent sRNA population^19^, confirming that sequence motifs can direct DNA methylation to gene targets. SR45 and other splicing factors have been linked to RdDM^77–80^, and related motifs may operate in both processes. RdDM dependence to enact epigenetic stress responses diminished significantly in advanced *msh1* memory generations (Fig. 1H, F), supporting a shift to maintenance methylation to sustain long-term heritable stress memory.

High-resolution and functional analyses identified a subset of 126 differentially methylated genes as putative regulators of environmentally responsive phenotype adjustment via pathways that trigger large-scale changes in alternative splicing behavior (Fig. 3, 5E). The network included spliceosome components and many targets of SR45 important for differential expression of isoforms. The largest nodes included *TOR*, *ILA*, *SYD* and other regulators of environmental stress response^51,53,54^, together with components of the RdDM pathway and chromatin remodelers. These hubs were alternatively spliced in some and differentially methylated in all *msh1* states. Hub genes were also enriched in the CTT motif, DMPs, and regulation by phosphorylation, a modification associated with alternative splicing and general response to environment^81,82^ (Fig. 5E).

It remains unclear how particular gbM repatterning leads to long-term adjustment in alternative splicing behavior. In linseed, repeated exposure to drought is co-regulated by alternative splicing and differential expression, while single exposure to drought causes only the latter^20^. In *msh1* memory, gbM can last for many generations and participate in adjusting growth phenotype. The site-specific nature of RdDM and its components, currently being investigated^19^, will likely reveal more direct connections to spliceosome components^77–80^. Thus, evidence for gbM influence on gene expression^11,20,24,26,32,33,70,71^ can no longer be ignored as an important factor in adaptive plant environmental responses.

## Methods

### Plant materials

All plant materials used in this study were *Arabidopsis thaliana,* accession Col-0. They were sown on pear mix in square pots, stratified at 4 °C for 48 hours and then transferred to a growth chamber at 22°C, 12 h light/12 h dark, and 120−150 μmol m^−2^ s^−1^ light intensity. *Msh1* memory lines were produced for Yang et al.^41^ and propagated through self-crossing for seven generations. Four gen1 non-memory lines (in addition to five screened by Yang et al.^41^) were screened for their memory potential in gen2 after self-crossing. The three sequenced gen2 non-memory individuals were screened for their memory potential in gen3 after self-crossing. The graft progeny line and msh1 mutant lines were derived previously in Kundariya et al.^34^.

### CRISPR-Cas9 lines

A binary vector on a pCAMBIA backbone was designed with a gRNA for *DMR2* exon 9 as described by Xing et al.^83^ using CRISPOR^84^. Memory gen 4 plants were transformed to obtain homozygous mutants by gen7 to compare phenotypes without the advanced generation of memory available. Col-0 was also transformed with both vectors and an empty vector was also transformed into memory plants to control for any transformation effects. Plants were watered as needed with 20-20-20 Grow More solution dissolved in deionized water at a concentration of 0.62g/L.

*Msh1* memory and Col-0 individuals were transformed using *Agrobacterium tumefaciens* containing a CRISPR-Cas9 construct. T1 seeds were surface sterilized with 20% bleach for 5 minutes and washed 5 times with sterile dH_2_O. They were plated on 0.5 MS 1% agar media supplemented with hygromycin (for positive selection) and cefoxitin (broad-spectrum contamination prevention). Positive transformants were transplanted to soil at the four-leaf stage and the growth protocol described above was followed. Putative mutants were confirmed in T2 and T3 by PCR amplifying the target sequence and Sanger sequencing (Extended Data Fig. 2). Phenotyping was done on several mutant lines in T3.

### Phenotyping

Plant photos were taken using an iPhone 12 camera at DAP as specified in figure captions. Rosette area was calculated in ImageJ by using the ‘Color threshold’ function. Bolting time was recorded for each plant as the days-after-planting (DAP). Rosette area was manually calculated in ImageJ by using the ‘Color threshold’ function.

### Sample processing for sequencing

Tissue for bisulfite sequencing and RNA-seq was collected from three biological replicates of memory gen7, new memory gen 1, and their full-sibling wild type looking plants. Above-ground rosette tissues was collected at bolting and flash-frozen using liquid nitrogen. Tissue was ground frozen and divided for DNA methylome and RNA sequencing. DNA was extracted using the DNeasy Plant Kit (Qiagen, Germany), according to manufacturer’s protocol. DNA yield was quantified Qubit Fluorometer with the high sensitivity dsDNA quantification kit (Thermo, Q32851) according to manufacterer’s protocol.

The remaining portion of ground tissue was used for sRNA extraction, using the NucleoSpin miRNA Plant Kit (cat#740971; Macherey-Nagel, Germany) following the manufacturer’s protocol.

For PacBio long read sequencing, leaf (at bolting) and flower (crown including open and closed buds) tissues were collected from memory, *msh1* mutant, graft progeny and WT control. RNA was extracting using NucleoSpin RNA Plant Kit (cat# 740949; Macherey-Nagel, Germany) following the manufacturer’s protocol, including DNase treatment step.

### High-throughput sequencing

Whole-genome bisulfite sequencing was conducted on the NovaSeq sequencing platform (Illumina, USA) at Novogene. 150bp paired-end reads were obtained with uniquely mapped reads and coverage listed in Table S2.

Small RNA sequencing was carried out at Huck Genomics Core Facility at Pennsylvania State University. The library was prepared using Illumina NextSeq 1000/2000 P1 reagents kit (cat# 20050264) according to manufacturer’s protocol and sequenced on Illumina NextSeq 2000 platform, generating 50 bp single-end reads.

For Iso-seq, RNA was converted to full length cDNA by reverse transcription with oligodT. The cDNA was cleaved by Uracil-Specific Excision Reagent (USER) enzyme and different transcripts were linked tandemly. Magnetic beads were used to select DNA fragments with size greater than 3.5kb that were amplified. SMRTbell adapters were ligated to cDNA and linear DNA without SMRTbell adapter. Library was qualified by QC. Sequencing was done on a PacBio SEQUEL II platform at BGI according to manufacturer’s protocol.

RNA-seq information differentiating memory from non-memory individuals was obtained by Yang et al.^41^ The msh1 translatome was obtained by Beltrán et al.^21^.

### Methylome analysis

Raw bisulfite sequencing reads were filtered for quality using FastQC. They were trimmed using TrimGalore with and Cutadapt. Then they were aligned to the TAIR10 reference genome using Bismark (version 0.23.1dev) with bowtie2.

Downstream differential methylation analysis was conducted as previously described ^38,45^ using the Methyl-IT suite version 0.3.2.6. (https://genomaths.github.io/methylit/). Briefly, context-specific Hellinger divergence (HD) was calculated between the methylation counts of the pooled control and all samples. For the analysis of memory methylomes, wild type or full siblings of memory with wild type phenotype were used as reference. Cytosines with a methylation level difference in control vs. treatment higher than the largest variation present in control samples (at least 20%), were selected. The selected cut-off point was evaluated using unsupervised machine learning to obtain true DMPs (α≤0.05, >95% classification accuracy) for each sample. Data for memory gen1 to gen6 obtained previously^41^ were re-analyzed, so all downstream analysis was based on the same version of the Methyl-IT software.

DMGs were classified with the count2test function as TAIR10 genes with at least 7 DMPs, a DMP density of 3 per 10^4^ with maximum coefficient of variance for each group at 1 (gen1 to gen6 with shallower sequencing) or DMP density of 3 per 10^3^ with maximum coefficient of variance for each group at 0.5 (remaining samples with deeper sequencing), log2fold changeLabove 1, and adjusted p-valueL<L0.05 according to the Benjamini and Hochberg method, and. DMPromoters were classified as 2kb upstream regions of TAIR10 genes, and DMTEs as all annotated transposable elements and pseudogenes in TAIR10, both with the same count2test criteria as DMGs.

### Discriminant analysis

The discriminant analysis was conducted with the *hclust* RStudio package stats (version 4.3.3) using Ward clustering including Ward’s clustering criterion (ward.2). The clustered statistic was total variation distance as calculated using Methyl-IT. This was done using all DMPs (Fig. 2b) or only DMPs within specific DMGs (Figs. 2c, d; 4a).

### Network analysis and ontology analysis

Gene network analysis was conducted using Cytoscape (version 3.10.3)^48^ as described previously^38,45^. The core hub network was obtained from DMGs present in all three msh1 states using the native k-means cluster function with Euclidian metrics, unconstrained number of clusters and 500 iterations. The node characteristics used to obtain the main cluster were betweenness centrality, closeness centrality, average shortest path length, clustering coefficient, degree, eccentricity, and topological coefficient.

For Fig. 3A, the resulting main cluster was arranged radially by their centrality using the yFiles tool in Cytoscpe. The size of resulting nodes was correlated with their degree and the transparency of their connections with the edge score. The division of nodes in the inner radius <2 (higher centrality) was added in Adobe Illustrator. The STRING Enrichment ^85^ option was used to add pie charts of enriched GO Biological process terms.

Enriched GO terms were identified using ShinyGO (version 0.80)^86^. The used cut-offs were false-discovery rate (FDR) < 0.05, p-value < 0.05 with redundancy removed. Extended Data Fig. 6 was plotted in RStudio using >5x fold significantly enriched Biological Process GO terms. Only terms containing >30 genes total are shown in the figure for legibility.

### DNA motif analysis

The sequence of 36 memory hub genes (Fig. 2C) and 67 msh1 hub genes (Extended Data Fig. 8A) was obtained from Araport11 and analyzed using Mutliple Em for Motif Elicitation tool from the MEME Suite (https://meme-suite.org/meme/tools/fimo)^87^ with default settings and any number of motif repetitions. As memory DMPs were primarily located in exons, exonic sequences were used to obtain the DNA motif, however, the genomic sequences also yielded the same motif with lower probabilities. To increase stringency in finding instances of the DNA motif in the genome, we searched for a 21-bp repetition of CTT in Araport 11 using Finding Individual Motif Occurrences (FIMO) from the MEME Suite.

### Small RNA analysis

Raw reads were trimmed and aligned using cutadapt (v2.8). To account for the non-canonical RdDM pathway, sRNAs of size 21bp to 24bp were included in the analysis. Trimmed reads were mapped to the TAIR10 genome using ShortStack the following settings: --bowtie_cores 8 --sort_mem 100G --dicermin 21 --dicermax 24 --foldsize 300 --mincov 20 --pad 100 --strand_cutoff 0.8. Samples were normalized using DEseq2 and differentially expressed sRNA clusters between different treatments were identified with the countTest2 function of the MethylIT suite. Only clusters differentially expressed between wild type and memory states were used for downstream analyses. Clusters with minimum of 5 reads per individual, |log2FC|L≥L0.5, α≤0.05 and false-discovery rate < 0.05, maximum coefficient of variance for each group of 1, and mapped to the nuclear genome were deemed significant. Overlaps of sRNA clusters and genomic features were identified using the findOverlaps function at different distances and filtered by unique hits. The percentages of overlaps in the figures were obtained by dividing the number of sRNA clusters within 1kb, 2kb, or overlapping genomic features by the total number of sRNA clusters for genes and TEs, and genomic features for DMG overlaps.

To quantify the PolIV- and CLASSY-dependence of the memory sRNA populations, we utilized the extensive datasets obtained by Zhou et al.^19^. All sRNA analysis was conducted after normalization with all samples from our analysis and GSE165001 ^19^ to allow valid comparison.

### Identification of differentially expressed isoforms

FLAIR v2.0.0 ^88^ was used to assemble and quantify isoforms. Sequence reads from PacBio SEQUEL II were aligned to the Arabidopsis genome (TAIR 10) using minimap2 (v2.26-r1175) with -ax splice:hq -t 40 -u f -G 6000 parameters (2022). The resulting bam files were sorted and non-aligned reads were filtered out. Bam files were converted to bed format using bam2Bed12 script provided as part of FLAIR. Misaligned splice sites were corrected using genome annotation (TAIR10 version 38 GTF) with FLAIR *correct* module and bed files as input using default parameters. FLAIR *collapse* was used to define high-confidence isoforms from corrected reads using default parameters. FLAIR *quantify* with default parameter was used to quantify isoforms in each sample. Finally, edgeR (v3.42.4) was used to identify differentially expressed isoforms between memory and WT control in flower and memory tissues. The reported alternatively spliced genes had at least one isoform differentially expressed between memory and wild type with |log2FC|L≥L0.5 and p-value < 0.05.

### Identification of alternative splicing event types

SUPPA (version 2.3.)^90^ was used to determine the event types in differentially expressed isoforms identified in the previous step. The function generateEvents --pool-genes with default parameters generated individual .gtf files for all event types and all samples. Events in individual isoforms could be separated into DCL-dependent and independent based on their presence in the *msh1* vs. *msh1, dcl2/3/4* comparison.

### Statistical analysis and visualization

The significance level of any overlaps between gene sets was calculated using the Hypergeometric test (phyper function in R or native function in Flaski heatmaps) with the total number of Arabidopsis genes set at 27,655.

All plots were made in RStudio using ggplot2 (3.4.4) and cowplot (1.1.3) and assembled into figures in Adobe Illustrator 2024.

The CTT motif and isoform presence was visualized using hierarchical clustering with Ward linkage in the Heatmap app of the Flaski toolset ^91^. The venn diagrams were constructed using the Venn diagram app of the Flaski toolset ^91^.

## Declarations

### Availability of data and materials

The next-generation sequencing datasets generated during the current study are available on the Gene Expression Omnibus (GEO) repository, under accessions GSE269287 (BS-seq), GSE269286 (Iso-seq), and GSE269527 (sRNA-seq). All remaining datasets generated and analysed during this study are included in this published article and its supplementary information files. The customized code used during the current study are available from the corresponding author on reasonable request.

### Authors’ contributions

AH, HK, and SAM designed the research. AH and HK performed the experiments, AUN participated in CRISPR mutant line creation. AH, HK, and RS participated in data analysis. AH made the figures. AH and SAM wrote the manuscript, with input from HK and RS. All authors read and approved the final manuscript.

## Acknowledgements

We thank John Howard, Seth McMahon, and Oliver Paulson for technical assistance. We thank Dr. Hong Ma for his critical feedback on the manuscript, and all members of the Mackenzie lab for valuable discussions.

