## Extended Data Fig. for "Intragenic methylation repatterning is associated with alternative splicing and unique epigenetic phenotypes"

**SUPPLEMENTARY DATA**

**Supplementary Data 1.** Methylome data bisulfite conversion rate and sequencing statistics, DMGs from memory gen1-gen7 (vs. wild-type), DMGs from new memory (vs. wild-type), DMGs from new memory (vs. wild-type-like full sibs), DMGs from memory gen1 (vs. wild-type-like full sibs).

**Supplementary Data 2**. Small RNA clusters in memory gen1 and gen7

**Supplementary Data 3.** Alternatively spliced genes and differentially expressed isoforms in three *msh1* states (vs. wild-type) and *msh1*, *dcl2/3/4* mutant (vs. *msh1*) in flower and leaf tissue (|log2FC| ≥ 0.5 and p-value < 0.05).

**Supplementary Data 4.** Gene descriptions of core hub DMGs identified in Fig. 3. The descriptions and locus names were obtained using the bulk download tool from The Arabidopsis Information Resource (TAIR)^1^. <https://v2.arabidopsis.org/tools/bulk/genes/index.jsp> on [www.arabidopsis.org](http://www.arabidopsis.org).

**Supplementary Data 5.** Number of phosphoproteins in *msh1* states gene sets and the significance of their overlap with *msh1* states’ gene sets.

| **Gene set** | **Number of genes in set** | **Number of phosphoprotein genes in set** | **% of phosphoprotein genes in set** | **Hypergeometric test p-value** |
| --- | --- | --- | --- | --- |
| All alternatively spliced genes in msh1 system | 8721 | 4285 | 49.2 | 6.06E-111 |
| All CTT genes | 9735 | 4677 | 48.0 | 1.37E-103 |
| DMGs (3 states) | 736 | 477 | 64.8 | 3.47E-45 |
| ASGs (flower) | 1526 | 907 | 59.4 | 1.94E-59 |
| ASGs (leaf) | 491 | 305 | 62.1 | 1.05E-24 |
| msh1 hub genes | 126 | 109 | 86.5 | 6.45E-28 |

**EXTENDED DATA**

**Legend**

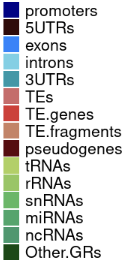

**Hypermethylated**

**Hypomethylated**

**CG**

**CHG**

**CHH**

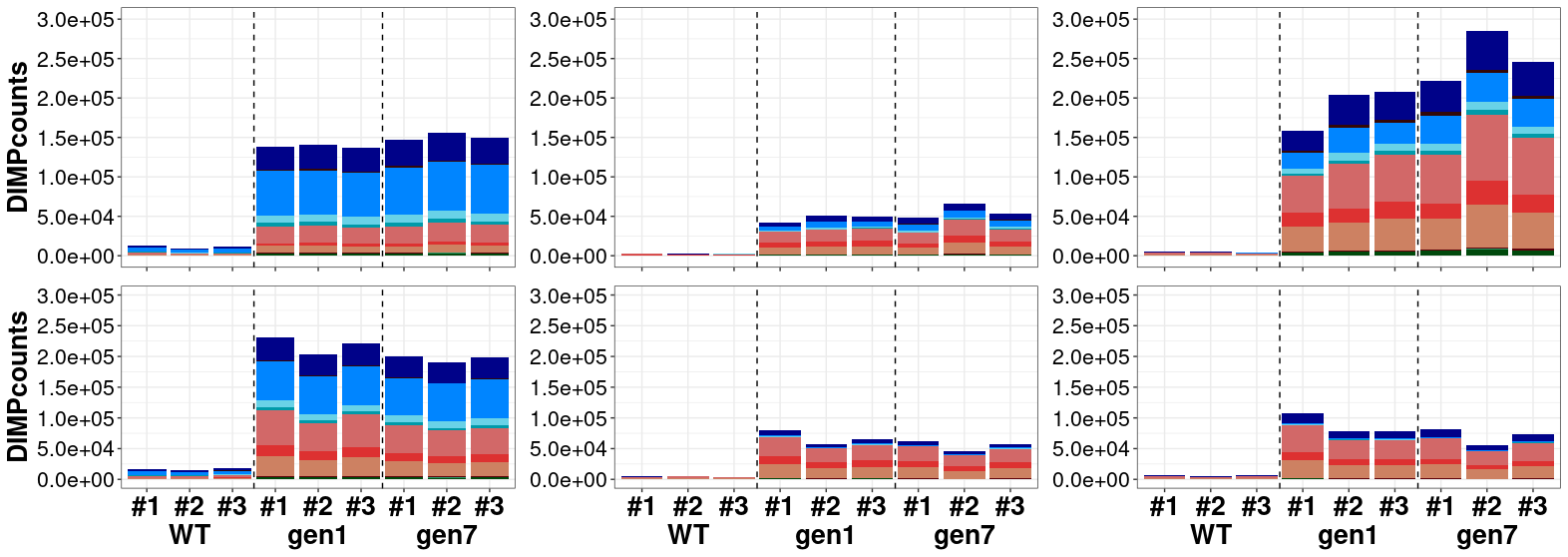

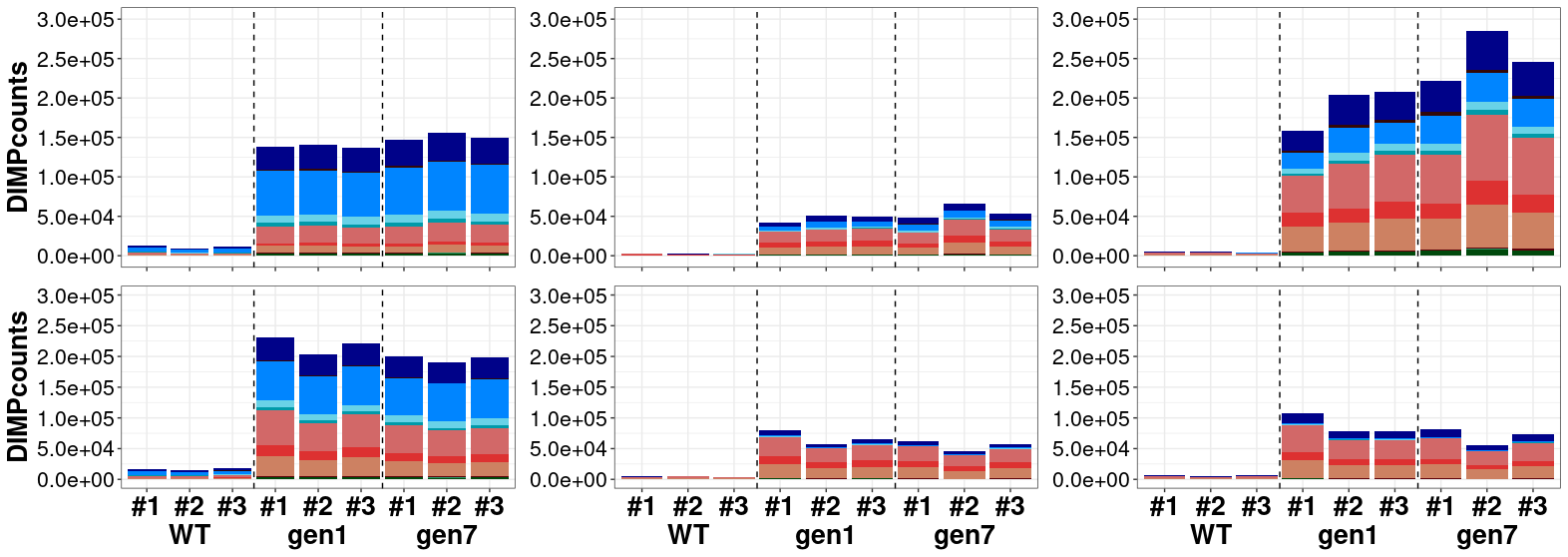

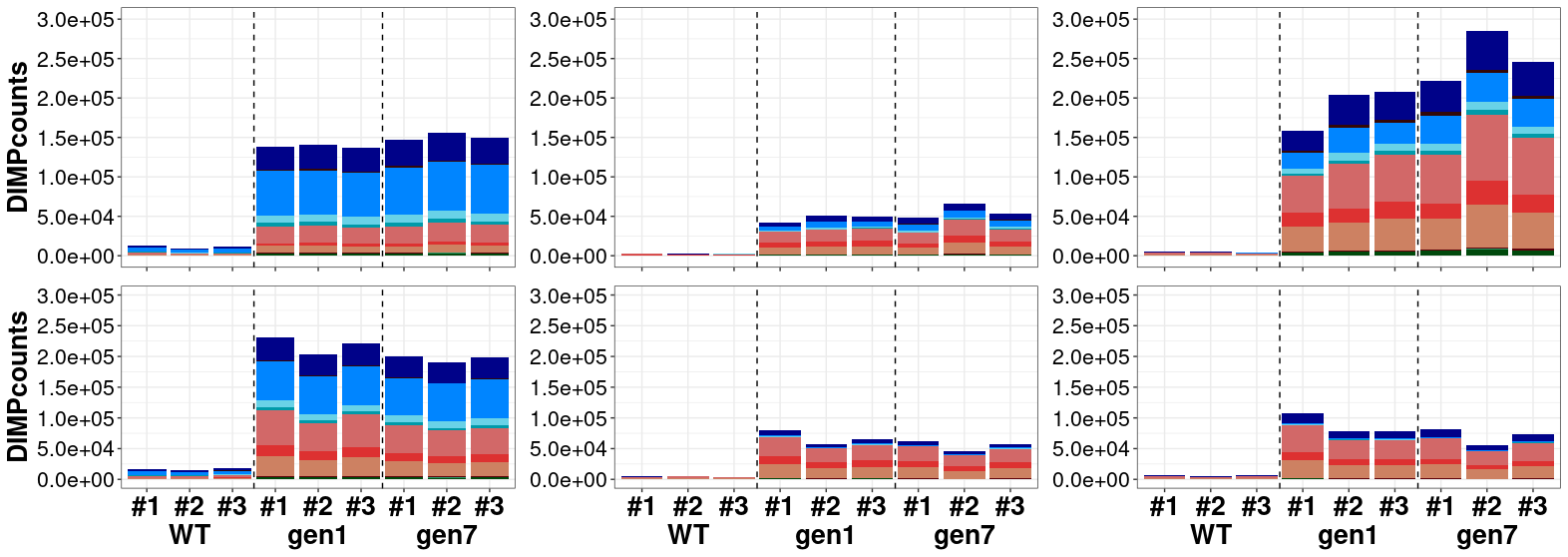

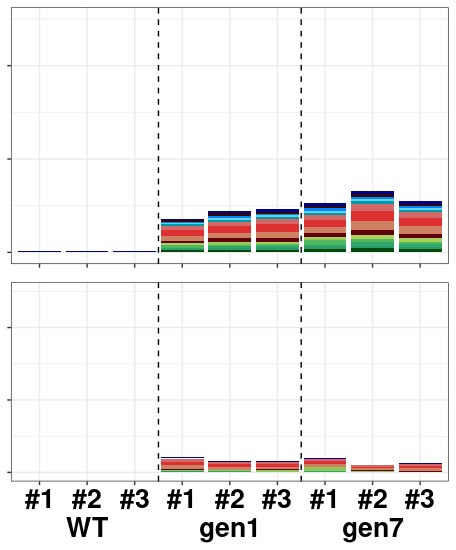

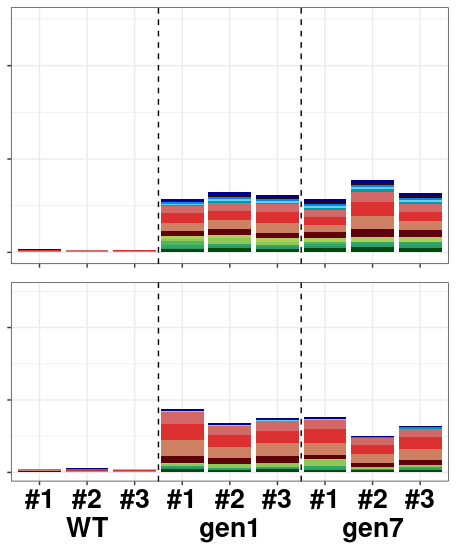

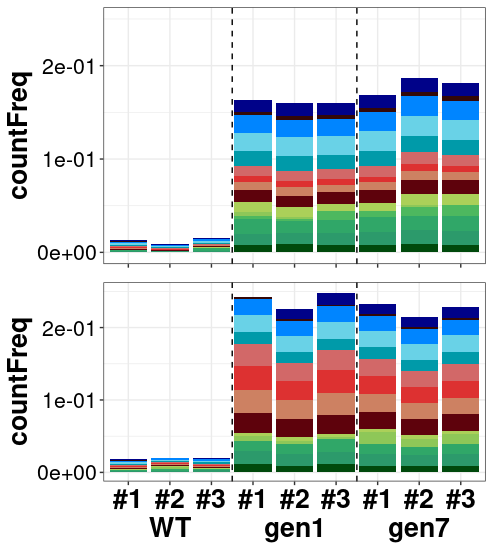

**B**

**A**

**Hypermethylated**

**Hypomethylated**

**Extended Data Figure 1. Memory is defined by gbM^T^ in the CG context. (A)** Differentially methylated positions (Hellinger divergence >20% control vs. treatment, GLM, α≤0.05, >95% machine learning classification accuracy) in each context and direction over three replicates of wild type (WT), new memory gen1 from non-memory, and gen7 inside genomic features and (**B**) normalized to the total number of potential sites for differential methylation in indicated contexts.

**Extended Data Figure 2. A drm2 mutation in memory does not affect the growth rate but alters bolting time.** (**A**) CRISPR mutations found in *DRM2* of memory gen7 and Col-0 visualized in Benchling. (**B**) Time to bolting recorded in days-after-planting (DAP) of in *drm2* mutants and control genotypes. After a significant Kruskal-Wallis test, all comparisons except mem gen7 drm2 CRISPR line1 vs. line 2 and mem gen7 vs. empty vector (n.s.) were significant at p<0.0001 with post-hoc pairwise Wilcoxon tests. (**C**) Growth curves of in *drm2* mutants and control genotypes. Each line represents a single plant. (**D**) Photos of in *drm2* mutants and control genotypes recorded 35 DAP.

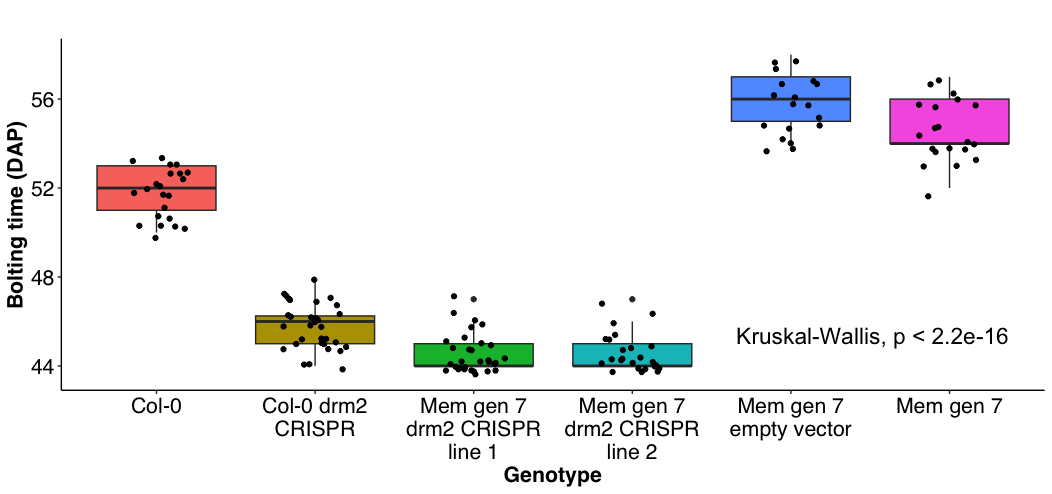

n.s.

n.s.

**B**

**C**

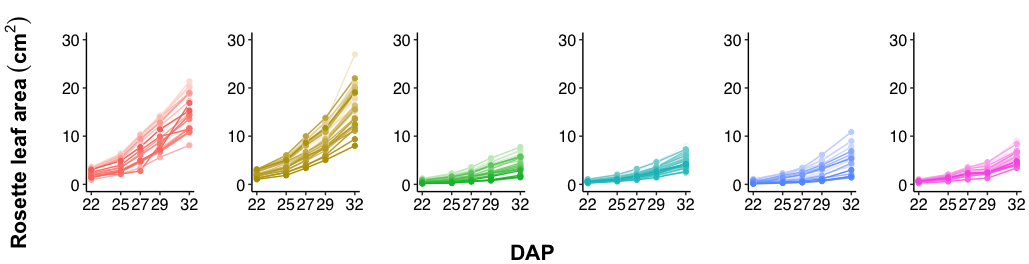

**D**

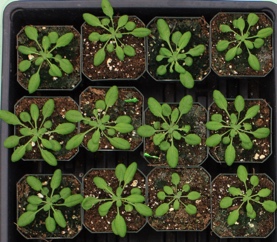

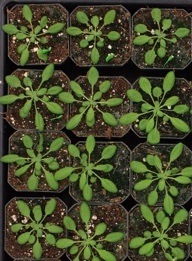

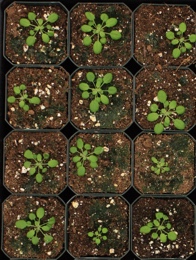

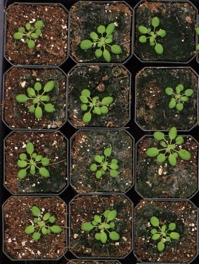

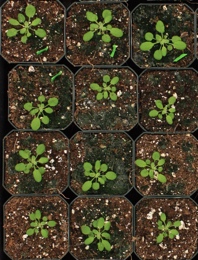

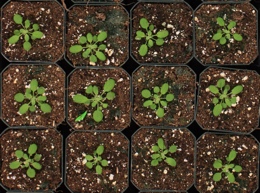

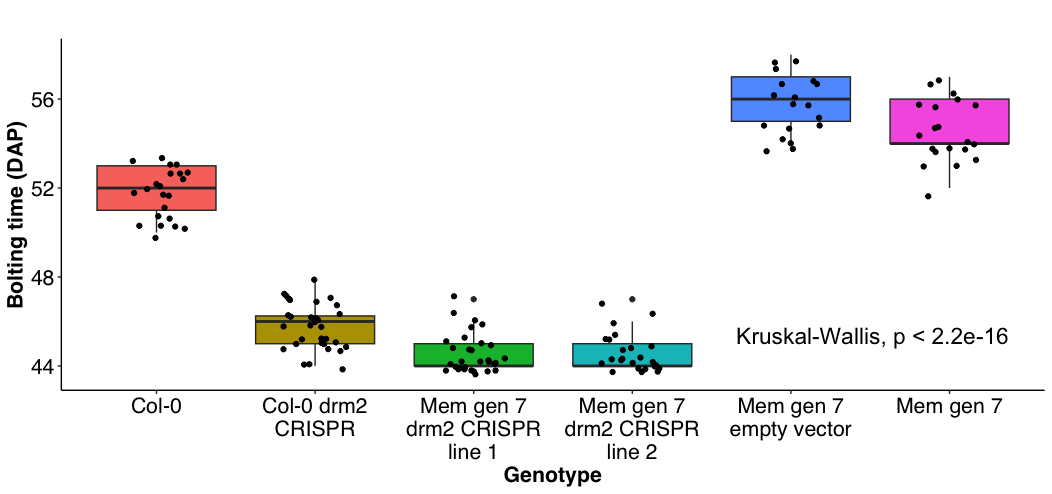

N=22

N=32

N=24

N=32

N=18

N=20

**A**

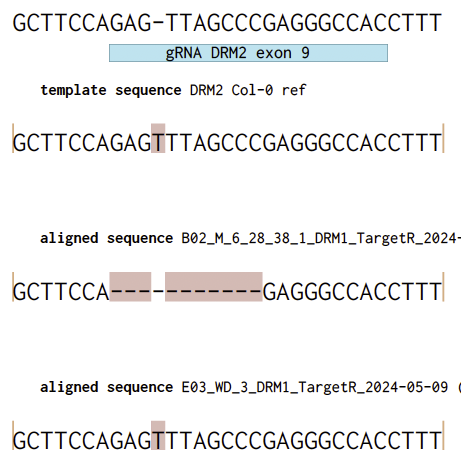

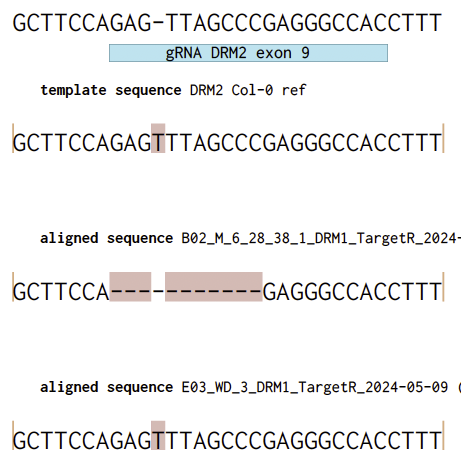

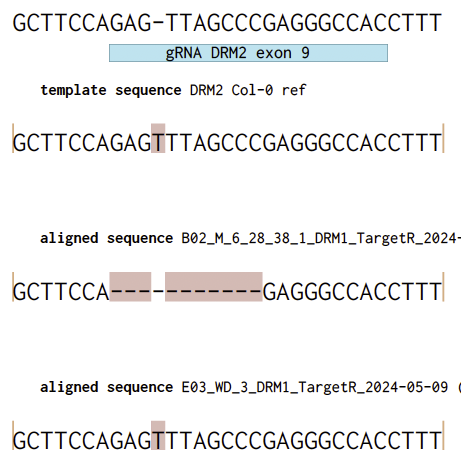

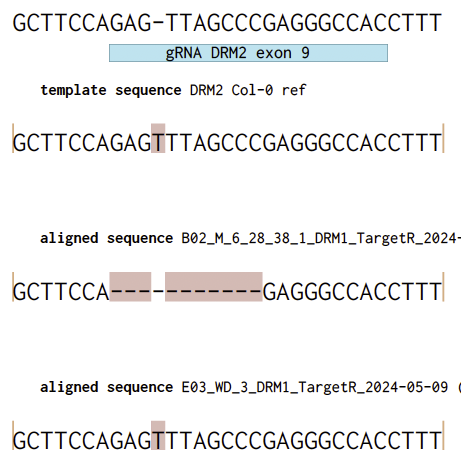

Col-0

Col-0 drm2 CRISPR

Mem gen7 CRISPR line 1

Mem gen7 CRISPR line 2

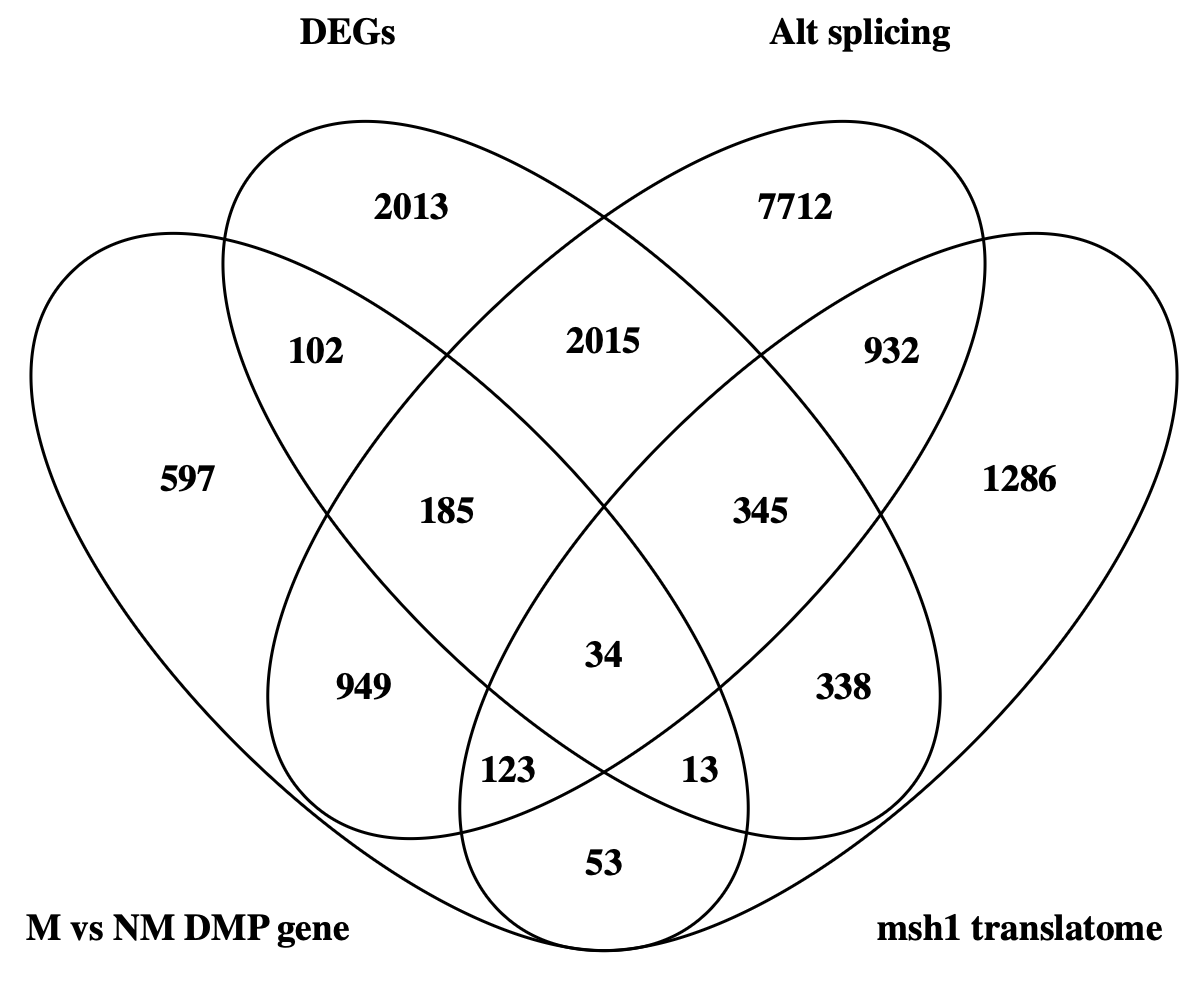

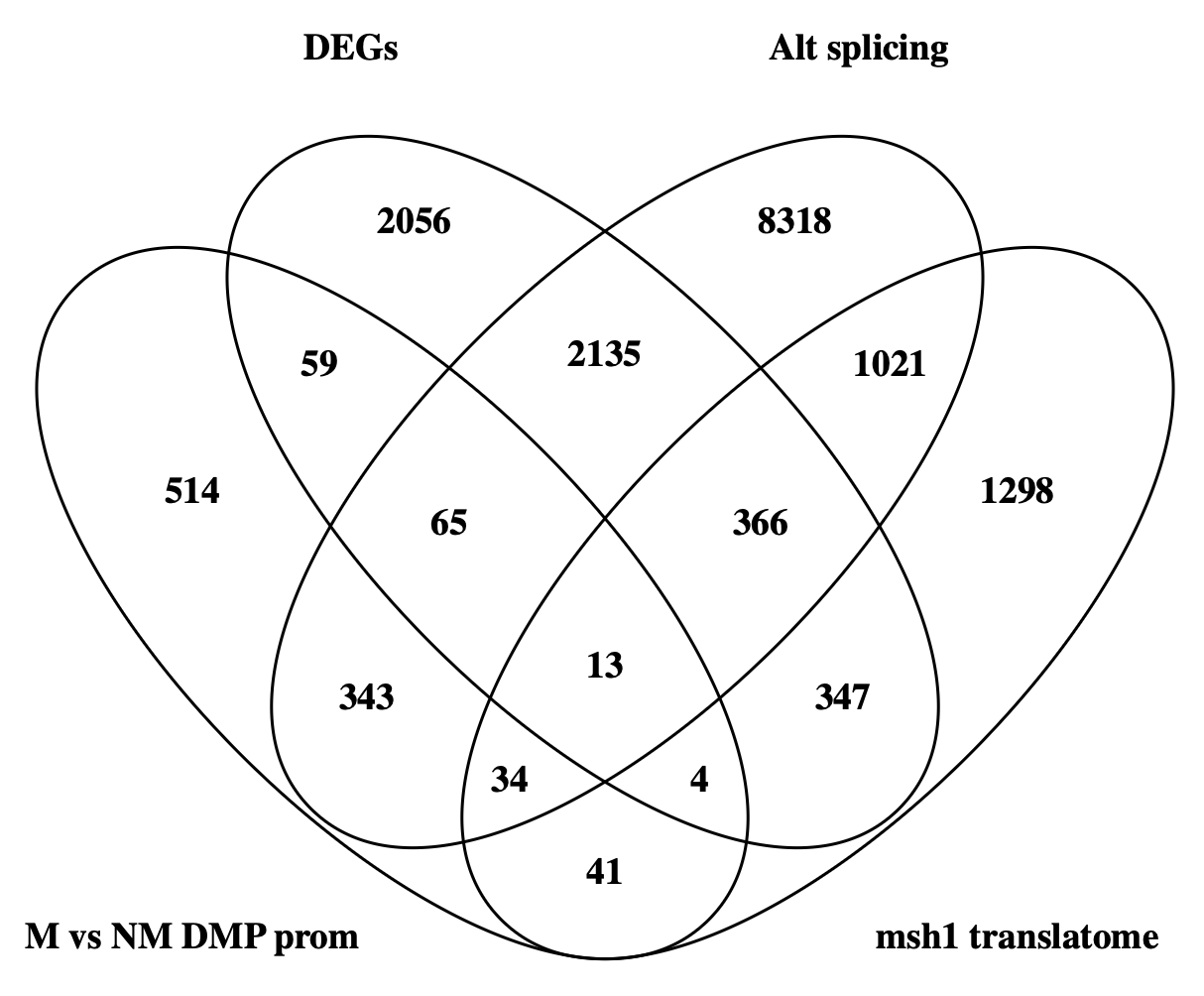

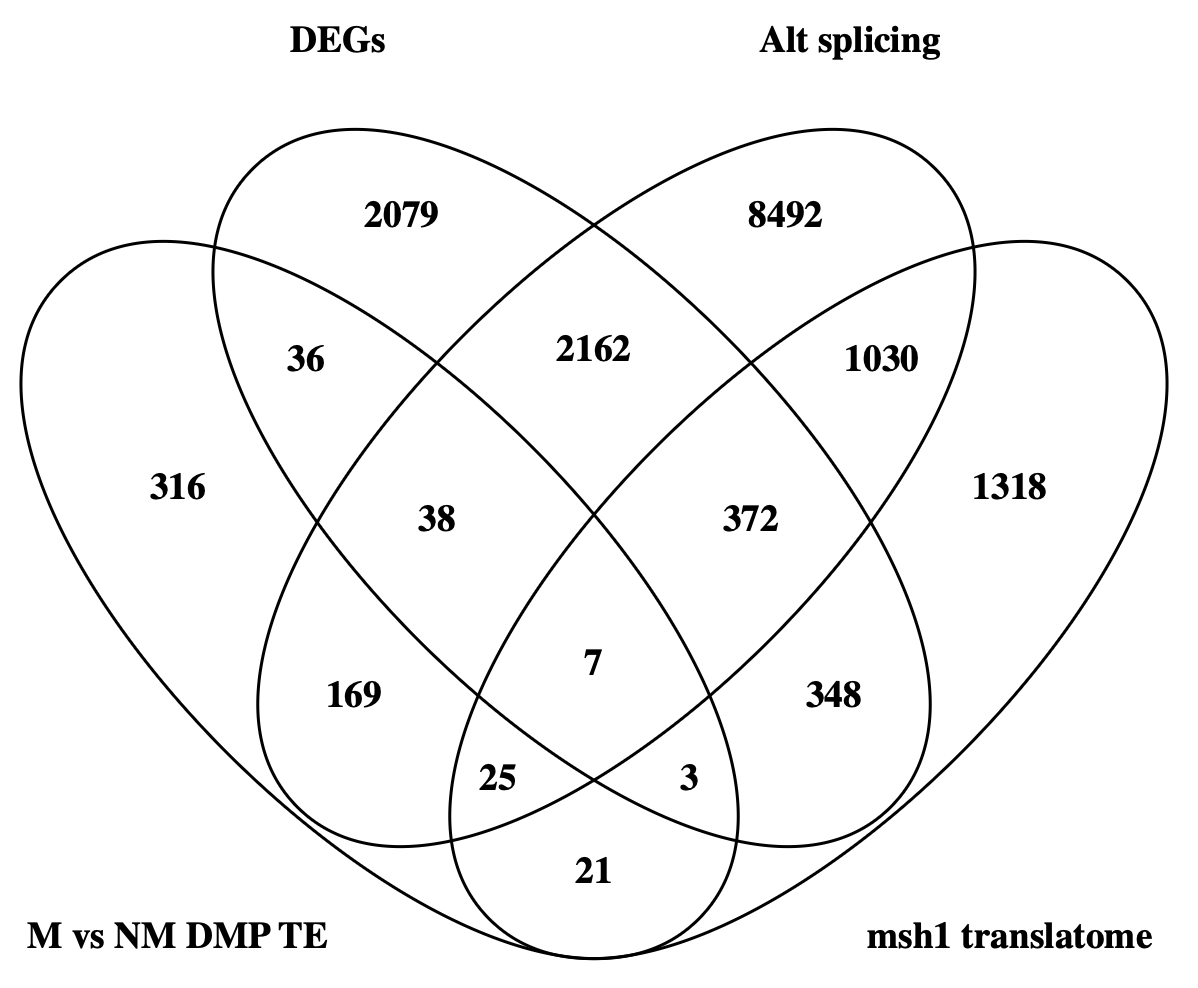

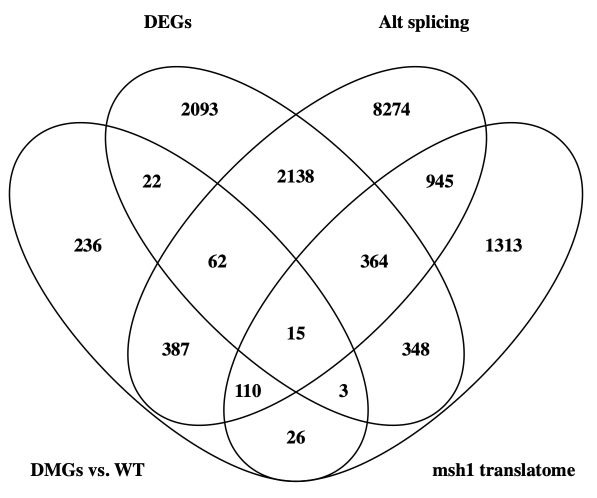

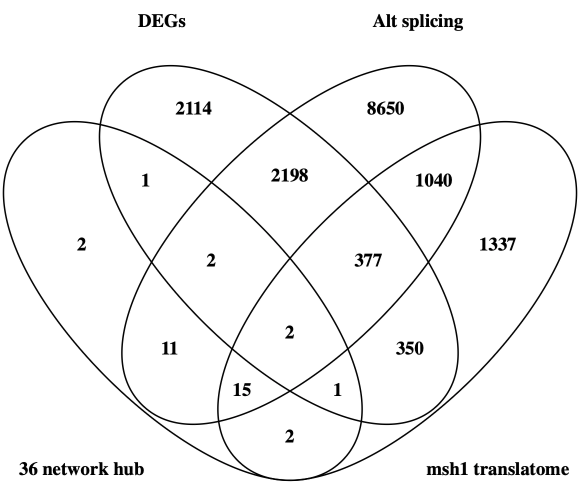

**A**

**B**

**C**

**D**

**E**

**Extended Data Figure 3. Memory differential expression is accounted for by differential methylation inside genes.** (**A**-**F**) Overlap between differential methylation in different genomic features and differentially expressed genes (DEGs), alternatively spliced genes (Alt splicing), and the msh1 translatome. Overlap of (A) methylation inside genes that differentiate memory and are inherited to gen 7 (M vs. NM DMP gene), (B) inside promoters (M vs. NM DMP prom), and (C) transposable elements (M vs. NM DMP TE) and differential expression. (D-F) Differentially methylated genes (DMGs) account for more differential expression than other differentially methylated features, which holds for (D) memory DMGs present in at least 6 of 7 generations (DMGs vs. WT), (E) and the 36-memory network hub.

**Extended Data Figure 4. The core hub memory genes are RdDM targets, but not in proximity to TEs.** (**A**-**C**) Core PPI network of 36 DMG, hubs discriminating memory from non-memory with potential. (A) 30/36 genes are overlapping sRNA clusters in memory gen1, which is reduced to13/36 in memory gen7. (B) According to *nrpd1* and *clsy1-4* sRNA clusters identified by Zhou et al.^2^, 36/36 of hub genes are PolIV-dependent and most are dependent on a CLASSY component. (C) 28/36 hub genes were identified as targets of *dcl2/3/4*-sensitive methylation by Kundariya et al.^3^ and 13/36 are within 2kb of a transposable element.

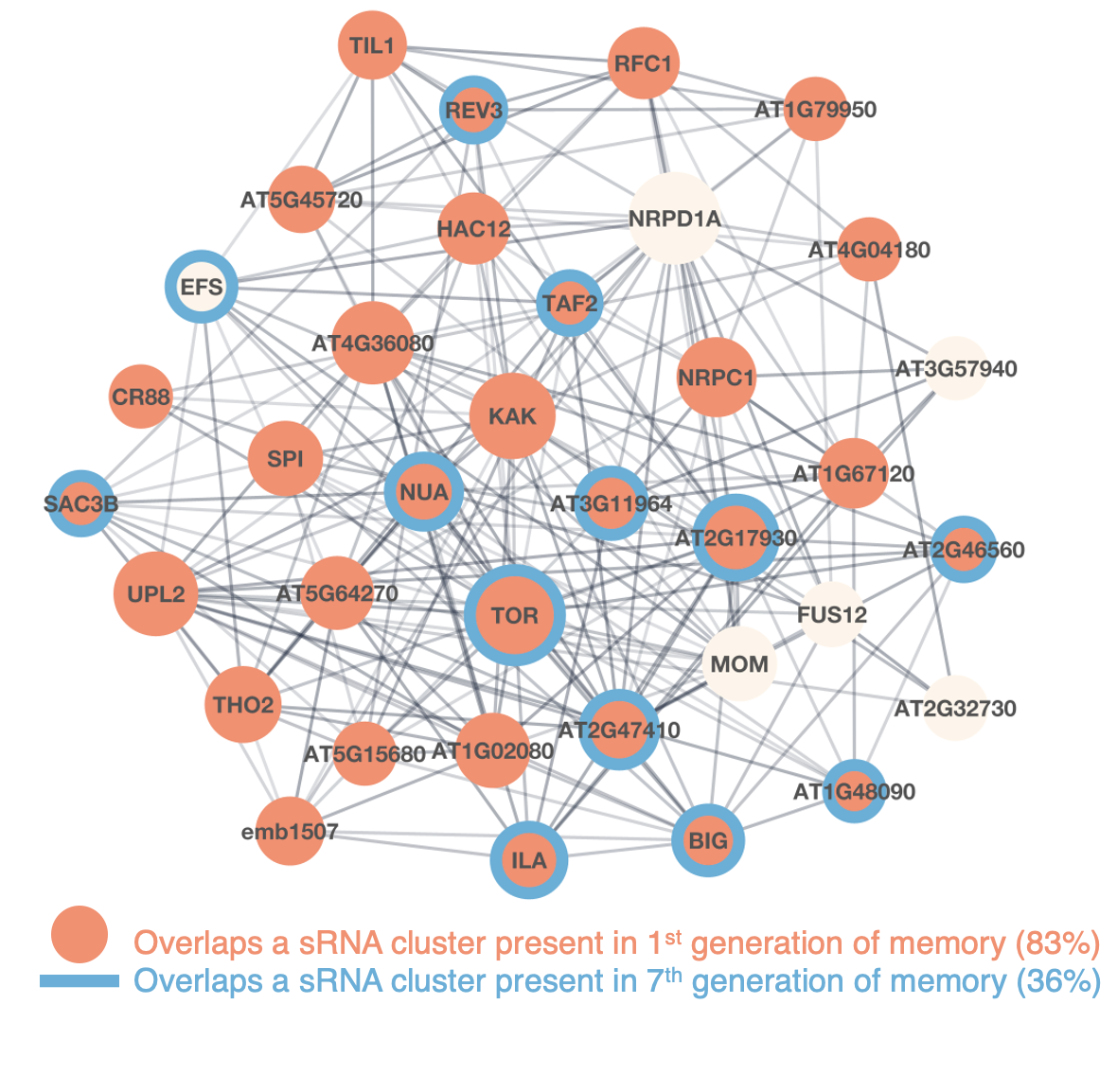

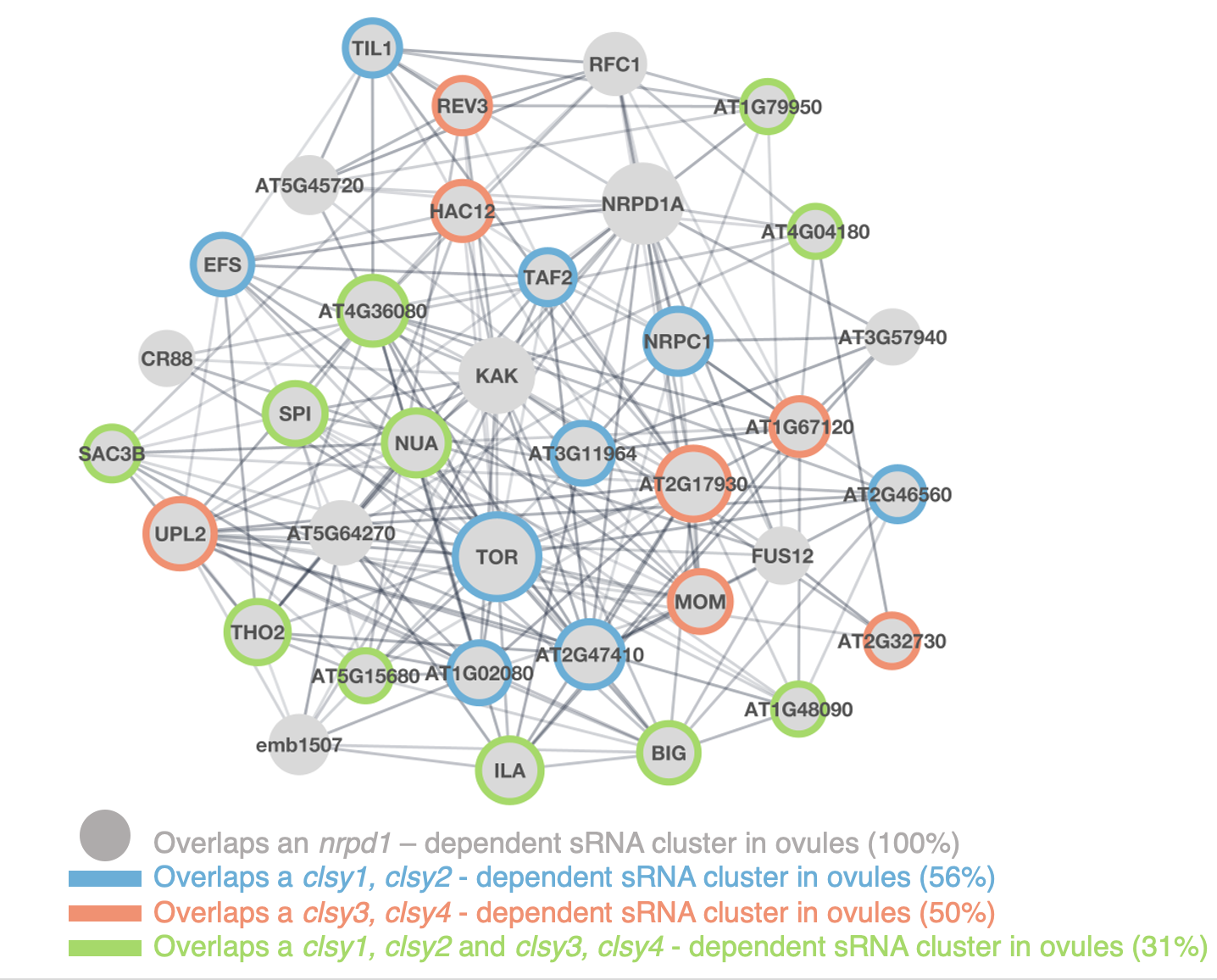

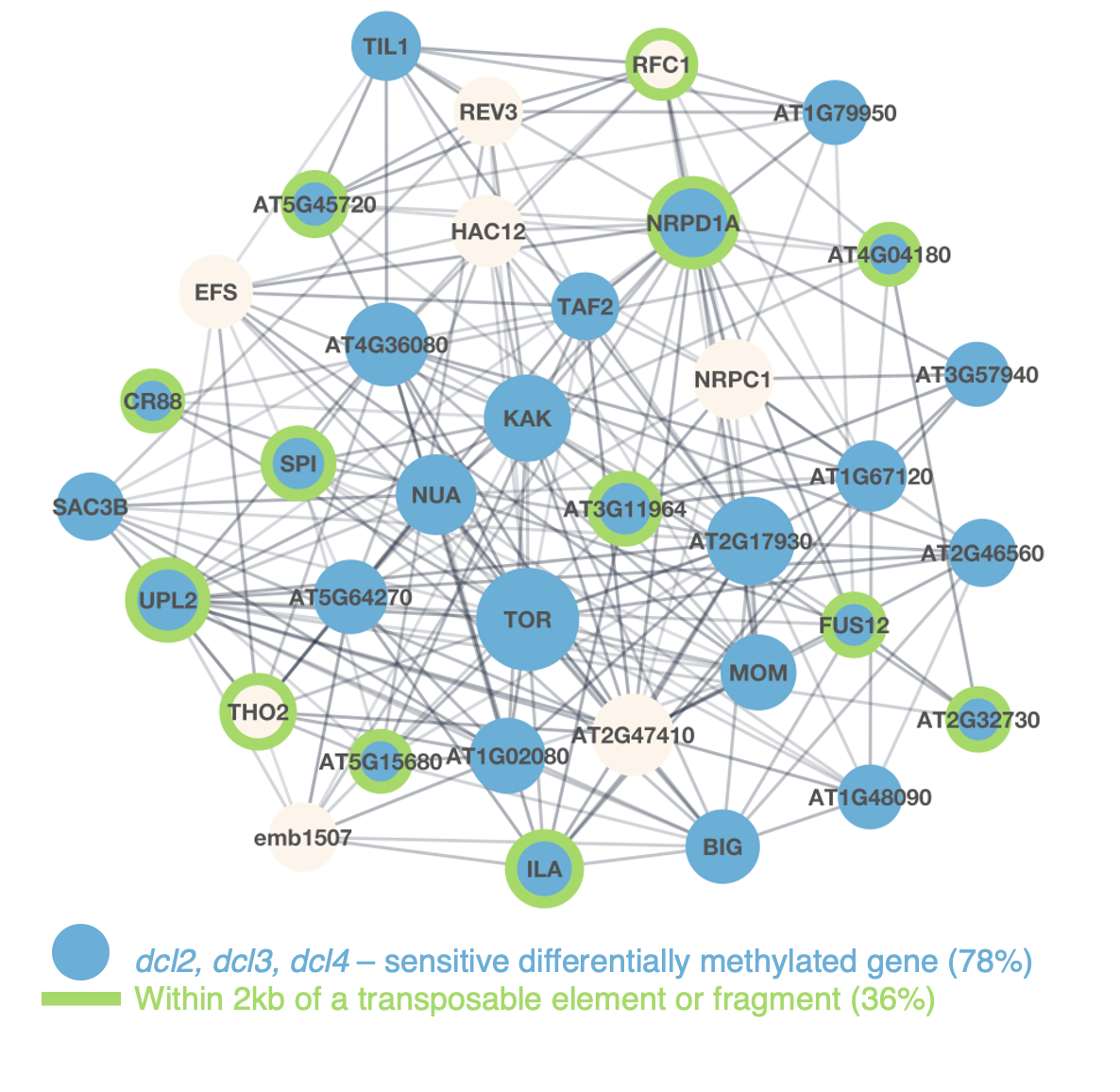

**A**

**B**

**C**

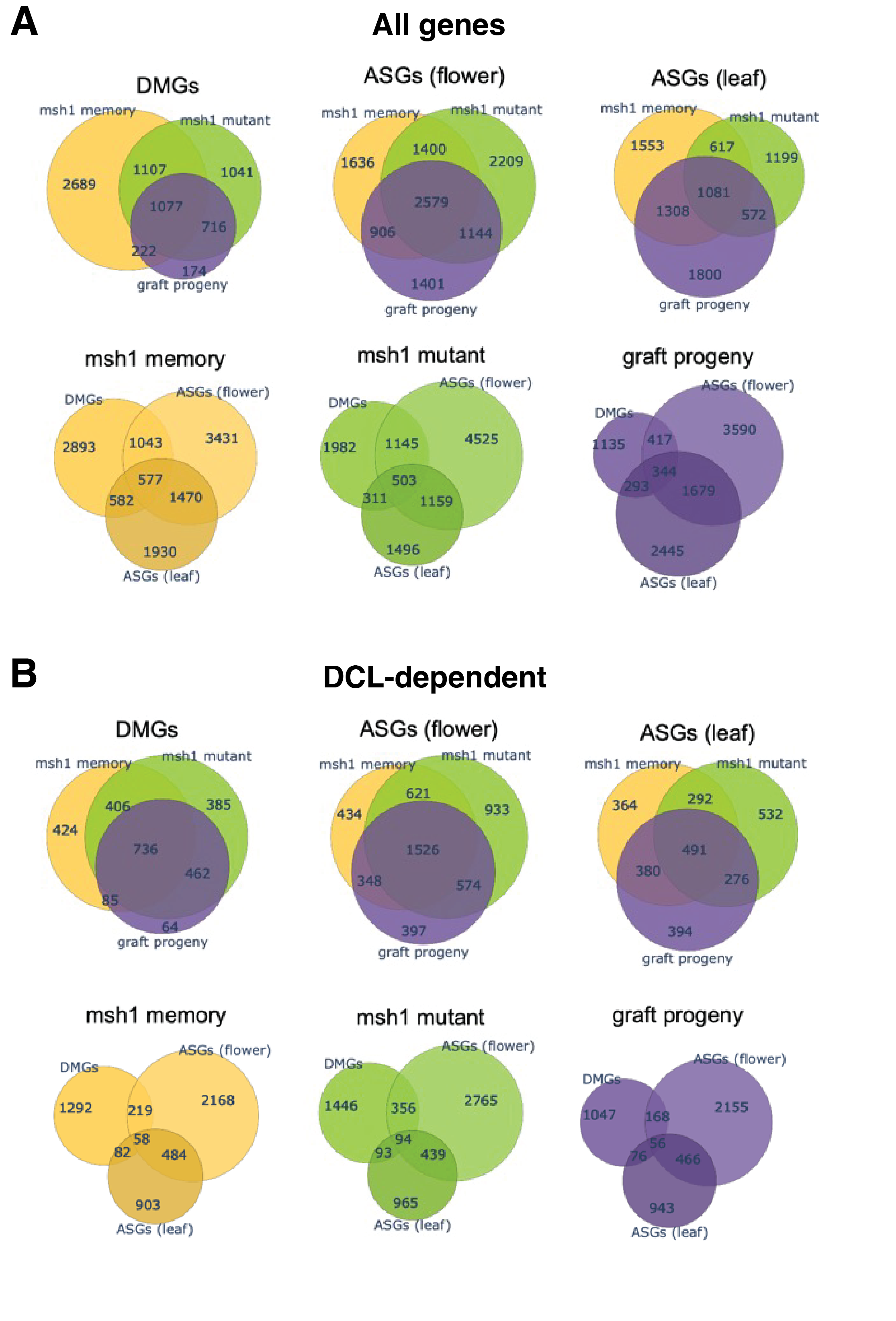

**Extended Data Figure 5. Overlap between alternatively spliced and differentially methylated genes.** (**A**) Venn diagrams showing the overlap of the three *msh1* epigenetic states (*msh1* mutant, *msh1* memory, graft progeny) in their alternatively spliced genes (ASGs) in leaf and flower tissues, and differentially methylated genes (DMGs). (**B**) Venn diagrams of gene sets shown in (A), subset to only show DCL-dependent genes, i.e. ones present in a comparison between *msh1* and *msh1, dcl2/3/4* mutants.

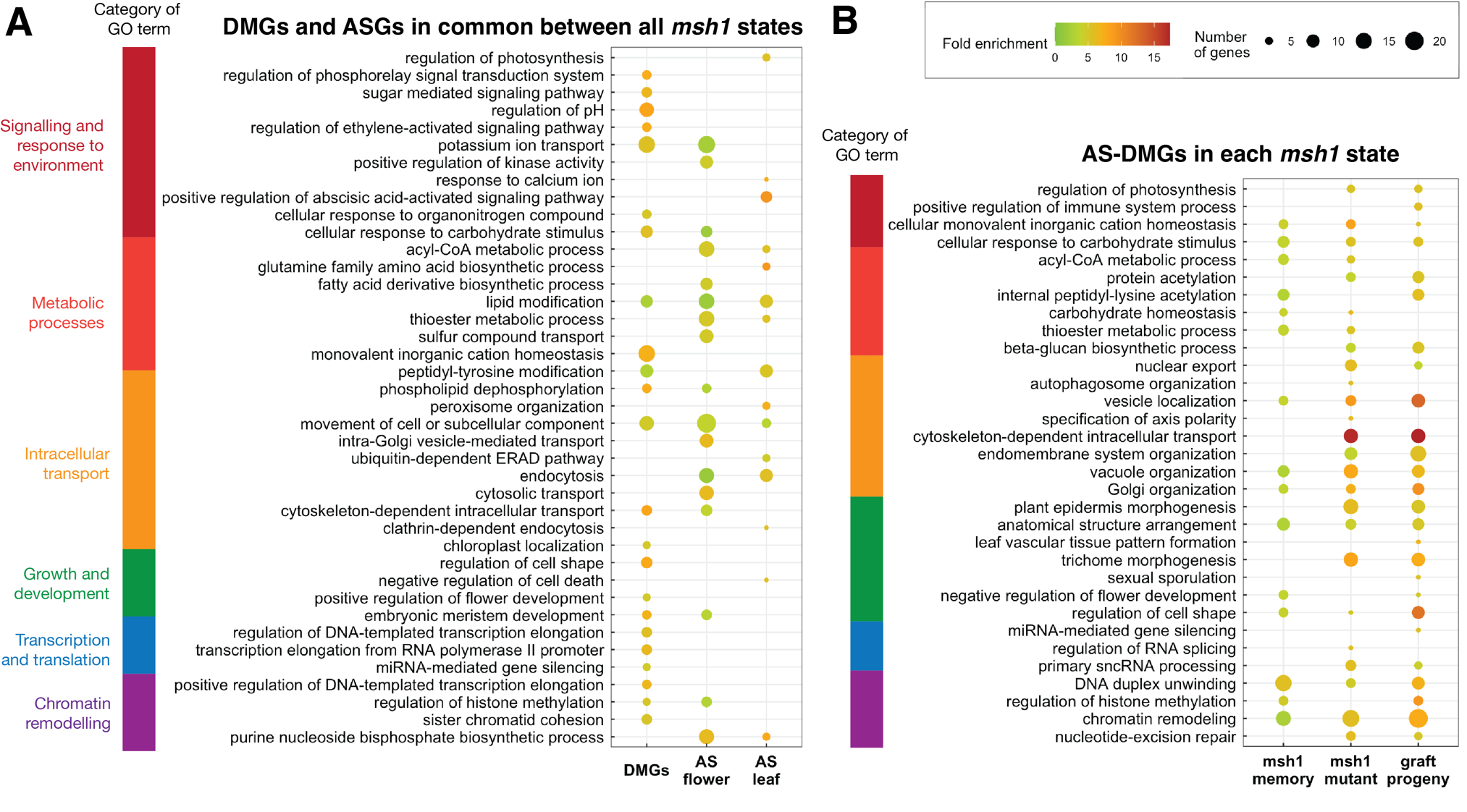

**Extended Data Figure 6. Differentially methylated and alternatively spliced genes are involved in distinct biological processes.** Enriched Biological Process GO terms (FDR < 0.05, p-value < 0.05, fold enrichment > 5x, total number of genes in GO term > 30) in (**A**) differentially methylated genes (DMGs) and alternatively spliced genes (ASGs) in flower and leaf tissue, present in all three *msh1* states and in (**B**) alternatively spliced and differentially methylated genes present in each of the three *msh1* states (*msh1* mutant, *msh1* memory, graft progeny).

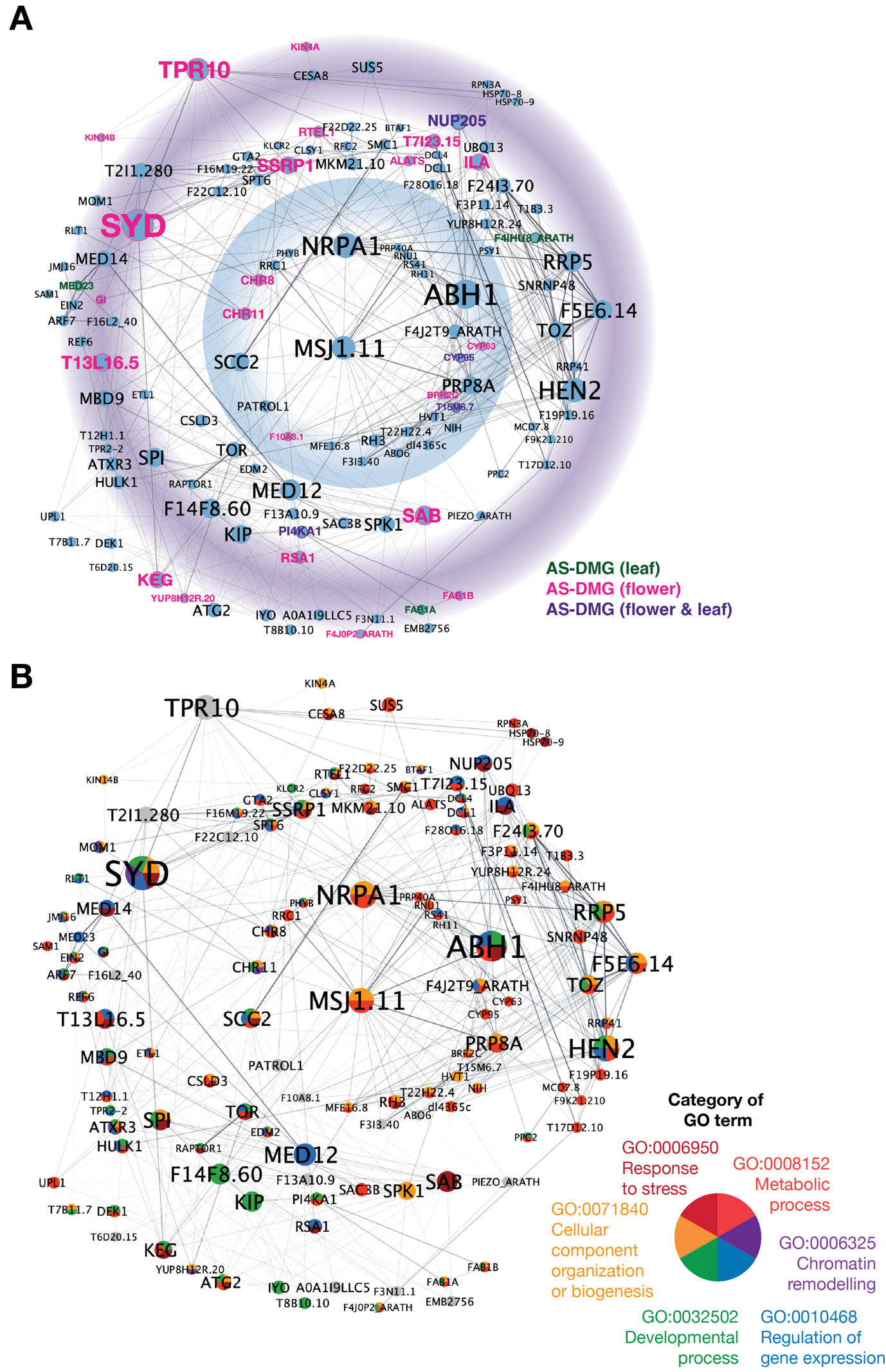

**Extended Data Figure 7. Core PPI network of 126 DMGs obtained using k-means clustering in Cytoscape from DMG common between all three *msh1* states.** Using the yFiles radial layout algorithm, the DMG nodes were ordered in concentric circles based on their centrality. The hub genes belonging to enriched GO terms as identified in Fig. S2 (FDR < 0.05) are colored according to the enriched GO term category.

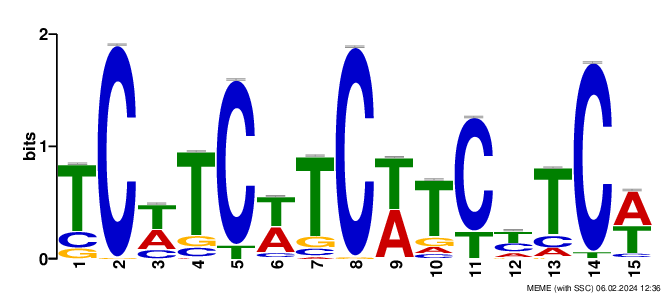

E-value=1.4e-072

**Extended Data Figure 8.** **Memory sRNA clusters overlap and exonic CTT motif.** (**A**) Logo plot of 15-bp motif identified within sequences of 67 core hub DMGs, identified by Kundariya et al. ^3^. The motif was identified using MEME (ungapped, 6-50 bp, any number of repetitions) and is present 335 times in the 67 hub genes but does not overlap with DMPs.

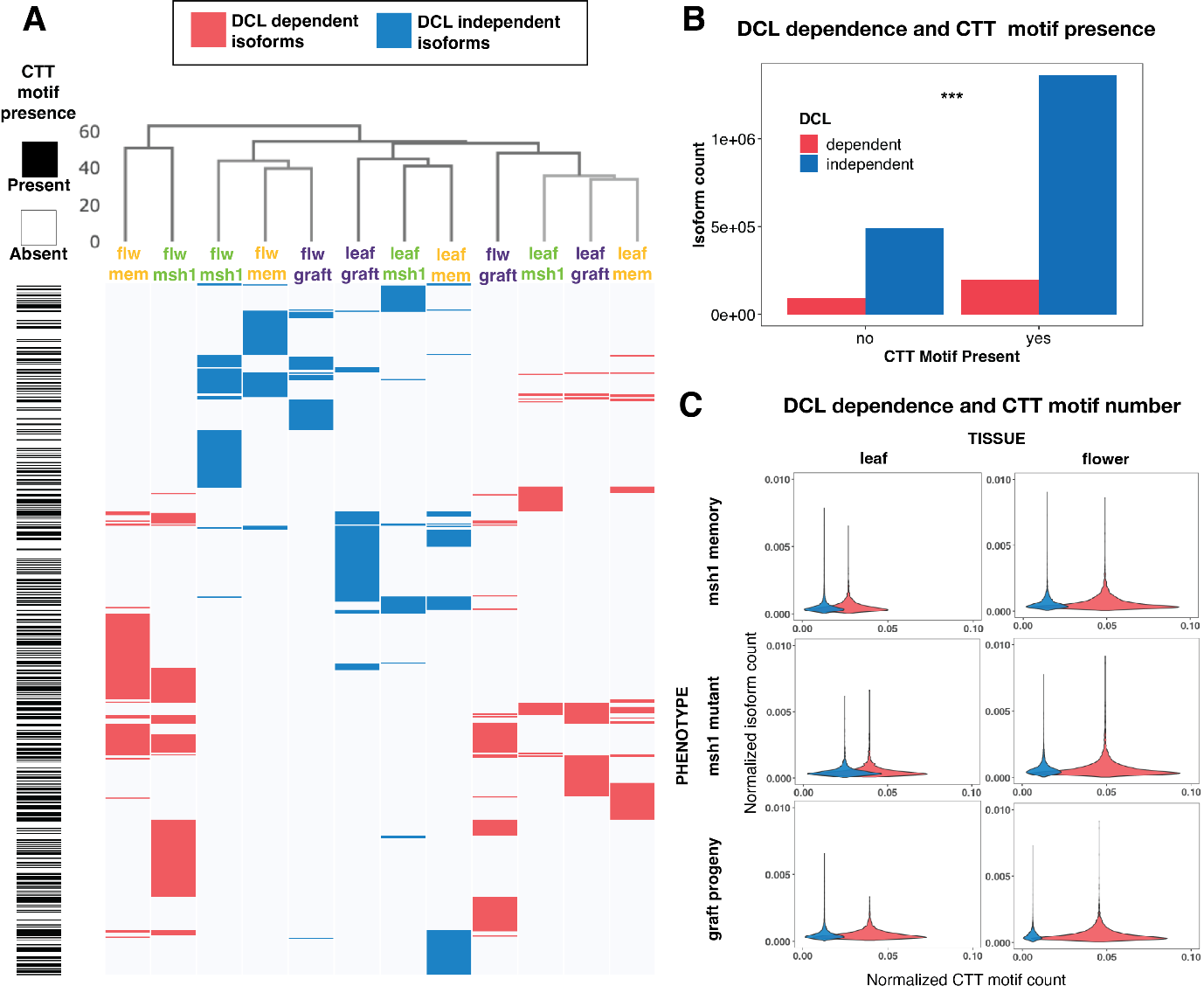
**Extended Data Figure 9. DICER-LIKE (DCL)-dependent alternatively spliced differentially methylated genes are associated with a higher number of CTT motifs.** (**A**) Hierarchical clustering of differentially expressed isoforms (|log2FC| ≥ 0.5 and p-value < 0.05) in *msh1* states’ leaf and flower tissue using Ward linkage (right). The presence or absence of a 21bp CTT motif is marked for each isoform (left). (**B**) Distribution of differentially expressed isoforms (|log2FC| ≥ 0.5 and p-value < 0.05) in *msh1* states’ according to the number of total isoforms in the gene of origin and the number of 21 bp CTT motifs overlapping it. Isoform are separated according to tissue (leaf and flower) and *msh1* state. (**C**) The presence of 21bp CTT motif inside DCL-dependent and independent (Pearson’s chi-squared, x^2^ = 3210.3, df = 1, pvalue < 2.2e-16)

**SUPPLEMENTARY INFORMATION**

**Supplementary Results**

**Alternatively spliced differentially methylated genes are core regulators of *msh1* states’ phenotypes**

To investigate DMGs with potential impact on phenotype through alternative splicing, we analyzed associated GO terms and DMG-, ASG-, and AS-DMG-derived gene networks. DMGs present in all three *msh1* states were enriched in GO Biological Process terms broadly associated with signaling and response to environment, chromatin remodeling, regulation of gene expression, and regulation of growth and development (Extended Data Fig. 6, 7). In contrast, ASGs in flower and to a lesser extent in leaf tissue were associated with GO Biological Processes in metabolic and intracellular trafficking categories (Extended Data Fig. 6A).

The three *msh1* phenotypes can be divided based on their epigenetic origins, with *msh1* and graft progeny derived from a direct RdDM-mediated connection to the loss of MSH1; the *msh1* memory state is seven generations removed from the initial downregulation of *MSH1*. Investigation of AS-DMGs present in each individual state revealed a division of enriched GO Biological Process terms consistent with this distinction (Extended Data Fig. 6B). While all three states were enriched in similar sets of environmental response and chromatin remodeling genes, the AS-DMG datasets associated with metabolism, intracellular trafficking, growth and development, and regulation of gene expression were distinct between the two phenotype groups (direct vs. transgenerational link to *MSH1* suppression). These observations suggested that the functions of genes targeted for differential methylation were distinct from those that were alternatively spliced, but AS-DMGs could enable the discrimination of epigenetic provenance of an *msh1* phenotype state.

To compare the function of genes in the core hub of *msh1* states with the larger set from which they derive, we searched for enriched GO Biological Process terms in the PPI network and sorted them in categories identified in Fig. 2 (Extended Data Fig. 7). Genes with the highest centrality were related to metabolic processes and organization of cellular components, followed by, both in frequency and centrality, regulators of gene expression, developmental processes and chromatin remodeling. Genes involved in stress response were found throughout the network. In addition to the GO terms present in Fig. 3B, RNA splicing (GO:0008380) was enriched in the network with 14 members, 13 of them found in the inner circle with highest centrality.

**Alternatively spliced and differentially methylated genes are regulated by phosphorylation**

In plants, phosphorylation regulates alternative splicing, and proteome studies have shown that splicing-related proteins are extensively phosphorylated^5^. To investigate if DMGs and ASGs in the *msh1* system are enriched in phosphoprotein genes, we utilized the PhosPhAt database of proteins regulated by phosphorylation ^6,7^. Phosphoproteins comprise approximately 39% of the Arabidopsis proteome. Of genes with differentially expressed isoforms in any of the *msh1* states, 49% coded for phosphoproteins (significant overlap, hypergeometric test, pvalue = 6.06E-111) (Supplementary Data 5). Similarly, of all genes containing the 21bp CTT motif, 48% were regulated by phosphorylation (pvalue = 1.37E-103) (Table S8). The fraction of phosphoproteins increased to 60-65% in ASGs in leaves and flower and DMGs present in all three *msh1* states (pvalue = 3.47E-45) (Supplementary Data 5). This enrichment for phosphoproteins was increased to 87% (significant overlap, hypergeometric test, pvalu6.45E-28) in the 126-gene hub of DMGs from Figure 3A (Supplementary Data 5). We interpret these data to indicate that key regulators of phenotype change regulated by differential methylation and alternative splicing are more likely to be phosphorylated, an indicator of their involvement in regulatory and signaling processes ^6^.
